# Integrated Constraint-Based Modeling of *E. coli* Cell-Free Protein Synthesis

**DOI:** 10.1101/2023.02.10.528035

**Authors:** Michael Vilkhovoy, Sruti Dammalapati, Sandra Vadhin, Abhinav Adhikari, Jeffrey D. Varner

**Author notes:** Corresponding author: Jeffrey D. Varner, Professor, Robert Frederick Smith School of Chemical and Biomolecular Engineering, 244 Olin Hall, Cornell University, Ithaca NY, 14853.

## Abstract

Cell-free protein expression has become a widely used research tool in systems and synthetic biology and a promising technology for protein biomanufacturing. Cell-free protein synthesis relies on *in-vitro* transcription and translation processes to produce a protein of interest. However, transcription and translation depend upon the operation of complex metabolic pathways for precursor and energy regeneration. Toward understanding the role of metabolism in a cell-free system, we developed a dynamic constraint-based simulation of protein production in the myTXTL *E. coli* cell-free system with and without electron transport chain inhibitors. Time-resolved absolute metabolite measurements for ℳ = 63 metabolites, along with absolute concentration measurements of the mRNA and protein abundance and measurements of enzyme activity, were integrated with kinetic and enzyme abundance information to simulate the time evolution of metabolic flux and protein production with and without inhibitors. The metabolic flux distribution estimated by the model, along with the experimental metabolite and enzyme activity data, suggested that the myTXTL cell-free system has an active central carbon metabolism with glutamate powering the TCA cycle. Further, the electron transport chain inhibitor studies suggested the presence of oxidative phosphorylation activity in the myTXTL cell-free system; the oxidative phosphorylation inhibitors provided biochemical evidence that myTXTL relied, at least partially, on oxidative phosphorylation to generate the energy required to sustain transcription and translation for a 16-hour batch reaction.

## Introduction

Cell-free systems are a widely used research tool in systems and synthetic biology and an emerging platform for biomanufacturing [1]. A distinctive feature of cell-free systems is the absence of cellular growth and maintenance demands, thereby allowing the direct allocation of carbon and energy resources toward a product of interest. Further, cell-free systems are amenable to direct observation and manipulation because they lack a cell wall, allowing rapid tuning of reaction conditions. The most widely used cell-free technology today is cell-free protein synthesis (CFPS). Combined with the rise of synthetic biology, cell-free protein systems have taken on a new role as a promising technology for just-in-time manufacturing of therapeutically important biologics and high-value small molecules. However, they have also been used for biosensing, prototyping genetic parts, metabolic engineering applications, and as educational tools at the high school and undergraduate levels for understanding synthetic biology [2, 3]. Commercial kits for different cell-free platforms, including *E. coli*, Chinese Hamster Ovary (CHO), HeLa, and plant cells [2–4], have accelerated the development of cell-free applications and technologies.

Cell-free protein synthesis systems have been used to explore fundamental biological mechanisms for decades. For example, cell-free protein synthesis systems in the 1950s and 1960s were used to understand transcription and translation processes. Borsook [5], Winnick [6], and Gale [7] used animal tissue homogenates and *Staphylococcus aureus* extracts to study amino acid incorporation into proteins, while the role of adenosine triphosphate (ATP) in protein production was explored by Hoagland [8]. Nirenberg and Matthaei demonstrated templated translation, i.e., the now familiar codon code, using *E. coli* cell-free extracts [9, 10]. On the other hand, Lederman and Zubay developed the first precursor to modern cell-free transcription and translation applications in 1967 when they developed a coupled transcription-translation bacterial extract using a DNA template [11]. More recently, the energy efficiency of *E. coli* CFPS was improved by generating ATP with substrate-level phosphorylation [12] and oxidative phosphorylation in the Cytomim system [13–15]. Since oxidative phosphorylation is a membrane-associated process, the study of Swartz and colleagues revealed that membrane-dependent energy metabolism can be activated in a cell-free system, suggesting complex metabolism is still occurring. Swartz and colleagues explored the scalability of cell-free systems, for example, for producing cytokines and antibodies on the industrial scale [16–18]. Commercially available platforms, such as myTXTL [19], use a different metabolic process that couples ATP regeneration and inorganic phosphate recycling to extend the duration of protein production. In addition, gene expression programs can be controlled using a variety of synthetic genetic circuitry [19, 20]. Thus, cell-free technologies are a viable alternative to living systems for investigative research, education, and small and large biomanufacturing.

Understanding cell-free systems’ metabolic performance is essential for optimizing their application [15]. A critical tool enabling this goal is mathematical modeling. Modeling the integration of cell-free transcription and translation processes with metabolism remains in its infancy, with few published mathematical models [21–23]. Horvath and coworkers developed an ensemble of dynamic *E. coli* CFPS models using kinetic parameter sets estimated from the metabolite and protein measurements [23]. This work expanded upon Wayman, and colleagues’ hybrid cell-free modeling approach, which integrated kinetic modeling with a rule-based description of allosteric control [24]. However, a critical challenge in the Horvath study was parameter estimation. Constraint-based approaches are an alternative strategy to fully kinetic modeling and the associated parameter estimation problem. Constraint-based models simulate metabolism under pseudo-steady-state conditions using constraints such as biochemical stoichiometry, and thermodynamic feasibility [25, 26]. Metabolic flux analysis (MFA) and flux balance analysis (FBA), which use stoichiometric reconstructions of metabolism, are examples of constraint-based approaches that have become standard tools in systems biology and metabolic engineering [27]. Stoichiometric reconstructions have been expanded to include the integration of metabolism with detailed descriptions of gene expression (ME-Model) [28–30] and protein structures (GEM-PRO) [31, 32]. Vilkhovoy and coworkers [22] developed an experimentally validated constraint-based model of CFPS, which integrated the expression of a protein product with the supply of metabolic precursors and energy. However, while the earlier Vilkhovoy model could reproduce (and in some cases predict) the expression of model proteins, several open questions were raised by the Vilkhovoy study.

In this study, we expanded on the previous sequence-specific constraint-based model of Vilkhovoy and coworkers [22] by integrating kinetic turnover rates, enzyme levels, kinetic descriptions of transcription/translation, enzyme activities, and absolute metabolite concentrations to constrain CFPS metabolic calculations. Metabolic flux estimated using flux balance analysis (FBA) and experimental inhibitor studies suggested that oxidative phosphorylation was active in the myTXTL system. Further, this activity was required to partially power protein production over the 16-hour cell-free reaction. Next, we explored an important mathematical question: the role of the objective function used in the flux balance analysis calculations. The previous study of Vilkhovoy and coworkers [22] assumed an objective of the maximization of the translation rate (which was also found to describe the data in the current study). However, in the cases of a perturbed network, e.g., protein production in the presence of inhibitors, optimal translation may no longer be a valid network objective. Toward this question, we used a complementary simulation approach, the Minimization of Metabolic Adjustments (MOMA), to predict the experimental data with and without inhibitors. Flux balance analysis and MOMA captured experimental mRNA and protein concentration data in the presence of electron transport chain inhibitors, with only minor differences between their predictions of metabolic flux distributions. In particular, MOMA predicted lower energy efficiency for transcription and translation than FBA, highlighting an increase in the wastage of energy resources in the treatment groups through unnecessary side reactions. Although small, the differences between the flux balance analysis and MOMA solutions could suggest that an optimal translation rate was no longer a valid objective function for the perturbed network.

## Results

### Integrated cell-free flux balance analysis (FBA)

Flux balance analysis is widely used in metabolic engineering to estimate intracellular metabolic flux. Flux balance analysis combines stoichiometric reconstructions of metabolism with a variety of experimental measurements, which constrain the flux calculation [33]. In particular, these measurements constrain the permissible bounds that a metabolic flux can have in the process of interest. Previously, Vilkhovoy and coworkers used flux balance analysis, in combination with time series metabolite measurements, to predict the synthesis of green fluorescent protein (GFP) and chloramphenicol acetyltransferase (CAT) under different promoters in different CFPS systems [22]. While the model predicted the overall protein synthesis rate well, there was significant uncertainty regarding the underlying metabolic flux distribution supporting protein expression.

To address this shortcoming, we developed a pipeline that integrated additional data, enzyme activity, and metabolite measurements into the cell-free flux balance analysis problem. Enzyme kinetic parameters, enzyme concentration, activity measurements, and metabolite concentration measurements were integrated into the metabolic flux calculation in the myTXTL system producing green fluorescent protein under a *σ*70 responsive promoter (Fig. 1). We adapted the earlier stoichiometric reconstruction of *E. coli* CFPS with 200 reactions (not including exchange reactions) and 157 metabolites which described glycolysis, the pentose phosphate pathway, tricarboxylic acid (TCA) cycle, amino acid biosynthesis, chorismate, purine, and pyrimidine metabolism, and transcription/translation processes, including tRNA charging and mRNA degradation [22]. Metabolic flux bounds were constrained by measurements (or associated literature values) using the enzymatic turnover and enzyme concentration measurements; turnover rates for each reaction were taken from BRENDA [34], or Adadi and coworkers [35]. Enzyme abundance was identified for 104 reactions in the myTXTL lysate by Garenne and coworkers [36]. Further, we validated enzyme concentrations for a subset of 15 reactions using enzyme activity assays. Four of the enzymes, hexokinase (*hk*), glutamate dehydrogenase (*gdh*), phosphoenolpyruvate carboxylase (*ppc*), and succinate dehydrogenase (*sdh*) were not reported by Garenne and coworkers. Still, they were found to be active in myTXTL with their corresponding activity assay. All remaining enzymes not reported by Garenne and coworkers were set to a median value of 50 nM. In addition, we constrained the upper bound of the corresponding reaction with the experimental enzyme activity level.

**Fig. 1:**
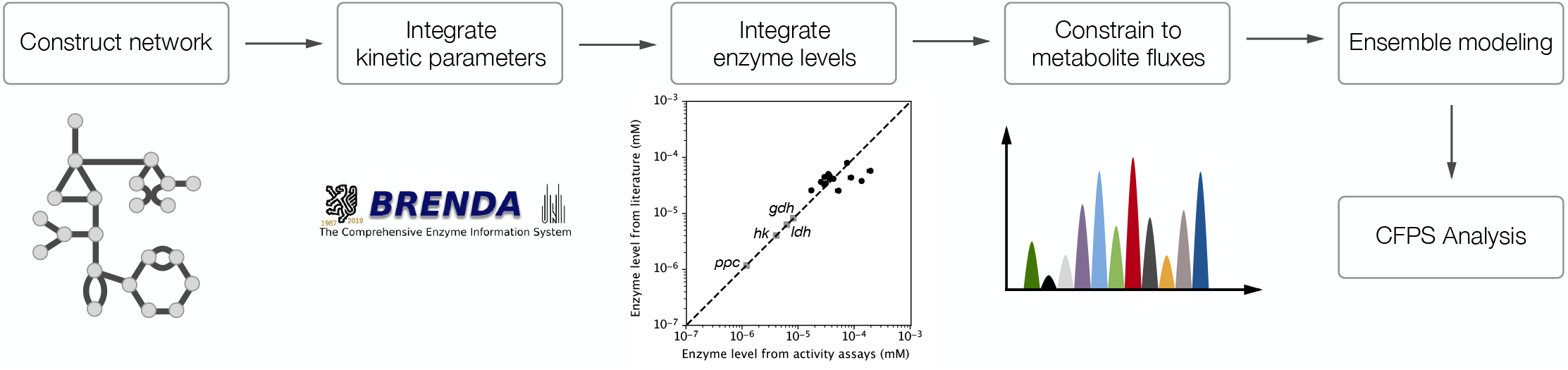
Schematic of the Integrated constraint-based modeling framework for cell-free protein synthesis. The metabolic network was adapted from Vilkhovoy and coworkers [22] where transcription/translation was integrated with metabolism. Maximum flux bound rates were formulated to be a function of the turnover rate and enzyme abundance found to be present in CFPS extract. Enzyme levels were validated for a subset of 15 reactions with enzyme activity assays. Four enzymes were not reported by Garenne and coworkers (grey boxes) [36]; these enzymes were found to be active with their corresponding enzyme activity assays. The metabolic flux for each time step was estimated while constrained to metabolic measurements where data was present (62 metabolites). Finally, the metabolic flux was sampled across an ensemble of 100 parameter sets constructed using experimental noise and literature parameters. Abbreviations: *hk* : hexokinase, *gdh*: glutamate dehydrogenase, *ppc*: phosphoenolpyruvate carboxylase, *sdh*: succinate dehydrogenase.

Absolute concentration measurements of ℳ = 63 metabolites, including central carbon, energy species, and amino acids, for the CFPS reaction time course, were also used to further constrain the flux estimation problem. Finally, the impact of uncertainty in measurements and model parameters taken from literature was addressed by simulating an ensemble of models (see materials and methods).

### Protein expression was dependent upon electron transport

Integrated flux balance analysis produced an estimated flux distribution that suggested oxidative phosphorylation and electron transport mechanisms were active during protein synthesis (Fig. 2). Metabolism in the myTXTL system has been reported to rely on maltodextrin and 3-phosphoglycerate (3PG) to provide energy resources for transcription and translation processes, with the involvement of oxidative phosphorylation being inconclusive [36, 37]. However, metabolic constraints and enzyme activity assays for glutamate dehydrogenase suggested that glutamate powers the TCA cycle and succinate dehydrogenase at least during the early time points. Flux calculations suggested oxidative phosphorylation was active throughout the CFPS reaction, with high flux at 2 hours (Fig. 2A) and a moderate flux at 8 hours (Fig. 2B). Maltodextrin and 3PG activated the glycolysis pathway, leading to an accumulation of organic acids such as pyruvate and acetate (Fig. S4). At 4 hours of the CFPS reaction, metabolism switched toward pyruvate consumption and valine synthesis, which accumulated in the extract (Fig. S2). Thus, analysis of the time-course CFPS flux distribution suggested the system relied on a mixture of metabolic processes to provide the necessary energy for transcription and translation. To probe the potential role of electron transport on protein production, the impact of electron transport inhibitors on cell-free metabolisms was measured.

**Fig. 2:**
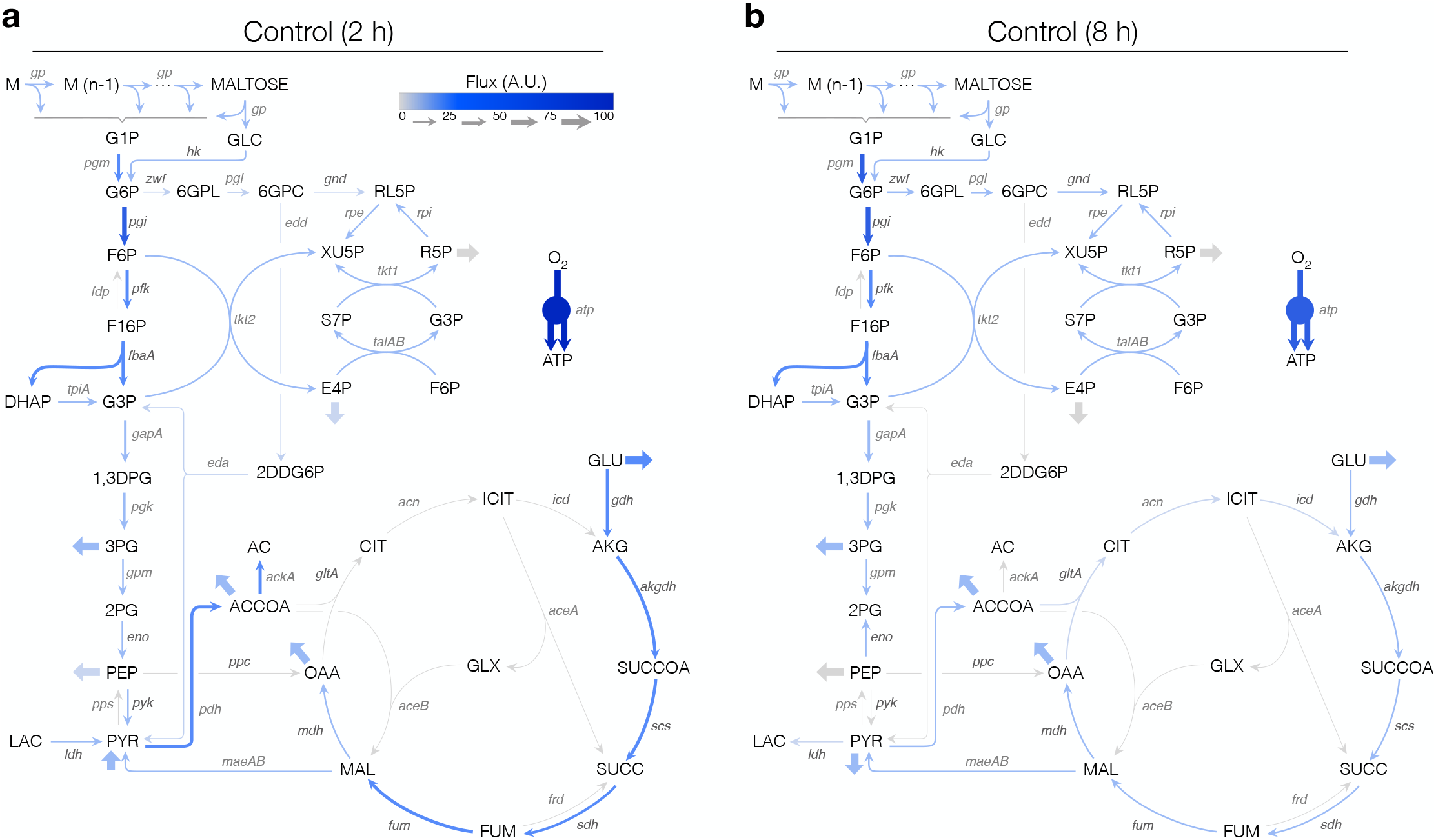
Mean flux distribution across an ensemble (N=100) for the control. Fluxes were determined by integrating kinetic parameters with enzyme levels and constraining to measurements of metabolites and enzyme activity levels where data was available. (a) Flux distribution at 2 hours of CFPS reaction. (b) Flux distribution at 8 hours of CFPS reaction. Fluxes were normalized to maltodextrin consumption at t=0 hours.

Inhibitor studies suggested that the transcription and translation of GFP depended on electron transport in the myTXTL CFPS system; mRNA and protein levels decreased in the presence of electron transport inhibitors (Fig. 3). The model described the dynamic behavior of mRNA and protein production for a nominal reaction (control) and reactions incubated with thenoyltrifluoroacetone (TTA) - an electron transport inhibitor of Complex II - or 2-4-dinitrophenol (DNP) - a membrane gradient uncoupling agent. The model captured the mRNA abundance for the control’s during the first 12 hours of the CFPS reaction; however, the model failed to capture the mRNA abundance at the 16-hour time point. In the simulation, CTP, GTP, and UTP were consumed toward the end of the CFPS reaction, transcription stopped, and mRNA was degraded. However, the experimental system maintained a non-zero steady-state mRNA concentration at 16 hours (Fig. 3A). Despite this, an model ensemble, considering uncertainty on the underlying model parameters, captured GFP production for the entire CFPS reaction (Fig. 3B). Thus, the sustained accumulation of GFP for the control compared to the electron transport inhibitors supported the hypothesis that oxidative phosphorylation was active in myTXTL.

**Fig. 3:**
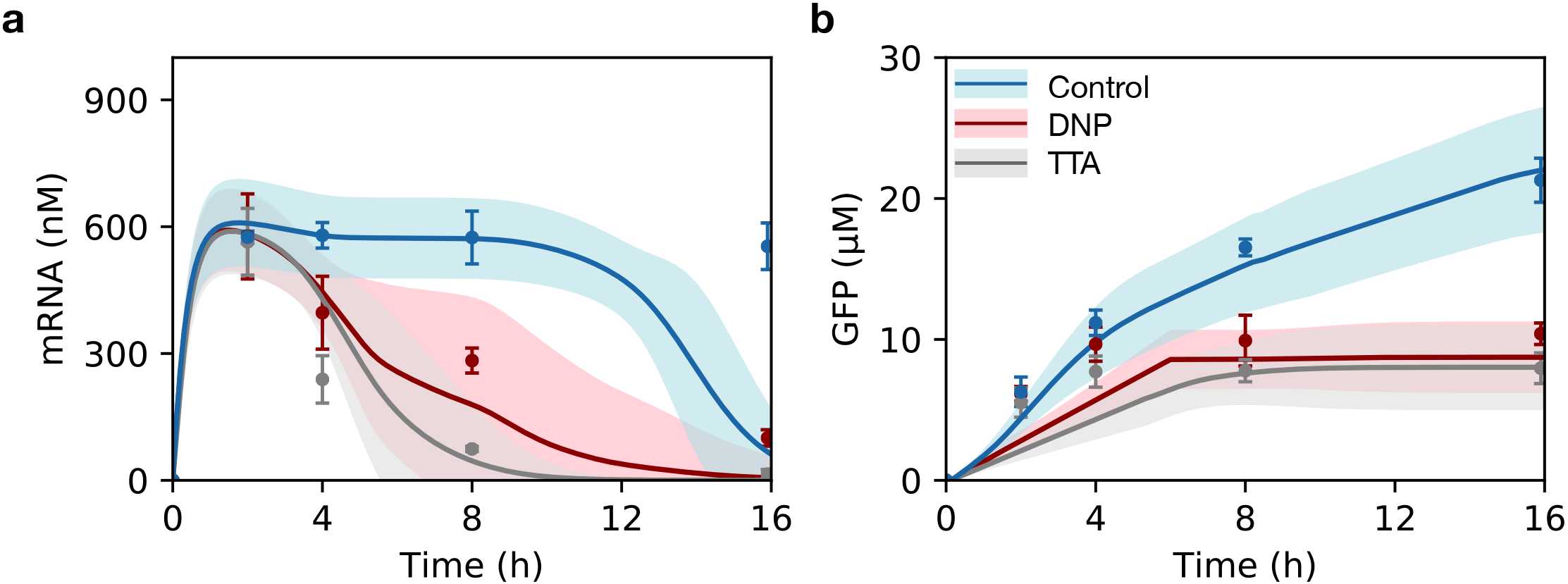
Prediction of mRNA and protein levels in CFPS for control (blue), DNP (red), and TTA (grey). The solid line denotes the mean of the ensemble (N=100), the shaded region indicates the 95% confidence interval, the points are experimental measurements, and the error bars represent the standard deviation of the experimental measurements (triplicate). The GFP mRNA and protein abundance was predicted, given the modeling framework for the three treatment cases.

When the CFPS reaction was incubated with DNP, oxidative phosphorylation was inactive at two hours (Fig. 4). At two hours, the model suggested 74% of the carbon traveled through glycolysis via *pgi* while 24% was diverted through the pentose phosphate pathway via *zwf*. The split toward pentose phosphate was notably higher for DNP-treated versus control reactions, where only 1% of the flux traveled through *zwf* at the two-hour mark of the reaction. The TCA cycle flux for DNP was similar to the control with high *gdh* activity across the ensemble. However, as the CFPS reaction progressed toward eight hours, significant differences appeared throughout the network. First, maltose was depleted, and thus lower *pgm* and *hk* fluxes were observed. In addition, 100% of the carbon traveled via *zwf*, and glycolysis’s first step ran backward to supplement G6P for the pentose phosphate pathway further. Lower glycolysis showed increased flux starting from *gapA* to *pdh* and towards *ackA*. Compared to the control, DNP had increased *gapA* and *pdh* flux. Finally, DNP treatment resulted in acetate accumulation, suggesting substrate-level phosphorylation was used to produce energy in the absence of electron transport processes.

**Fig. 4:**
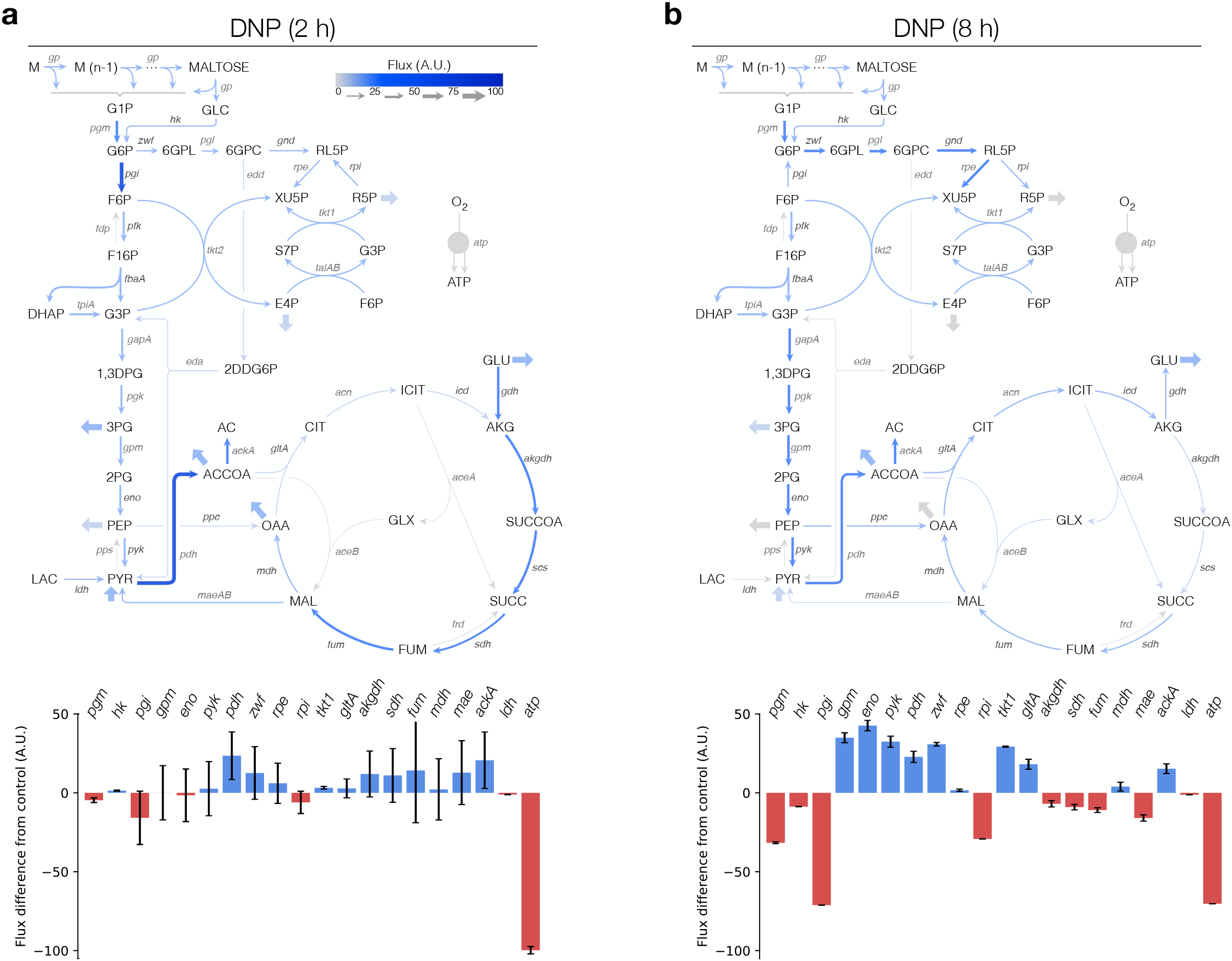
Mean flux distribution across an ensemble (N = 100) for DNP treatment. Fluxes were determined by integrating kinetic parameters with enzyme levels and constraining to measurements of metabolites and enzyme activity levels where data was available. (a) Flux distribution at 2 hours of CFPS reaction. Flux differences from control are shown at t = 2 hours of CFPS reaction. (b) Flux distribution at 8 hours of CFPS reaction. Flux difference from control shown for key reactions at 8 hours of CFPS reaction. Fluxes were normalized to maltodextrin consumption at t = 0 hours.

When the CFPS reaction was incubated with TTA, the rate of oxidative phosphorylation decreased but remained active (Fig. 5). Upper glycolysis involving maltodextrin consumption, glucose-1-phosphate, and glucose utilization was similar when compared to the control with a slight increase in *pgm* activity. However, just as in the case with DNP, there was a higher split toward the pentose phosphate pathway, with 15% at 2 hours of the reaction and 18% at 8 hours of the reaction. The most notable differences were in the TCA cycle which had lower activity, which was further supported by high glutamate levels in the media. TTA is an inhibitor of succinate dehydrogenase and uncoupled the TCA cycle from central carbon metabolism. In the presence of TTA, oxidative phosphorylation remained active, but at a 51% reduction when compared to the control. Lastly, despite having an active oxidative phosphorylation reaction, central carbon metabolism showed significantly less flux compared to both the control and DNP. In addition, there was a higher accumulation of lactate with approximately 8 mM at the end of the 16 hour reaction.

**Fig. 5:**
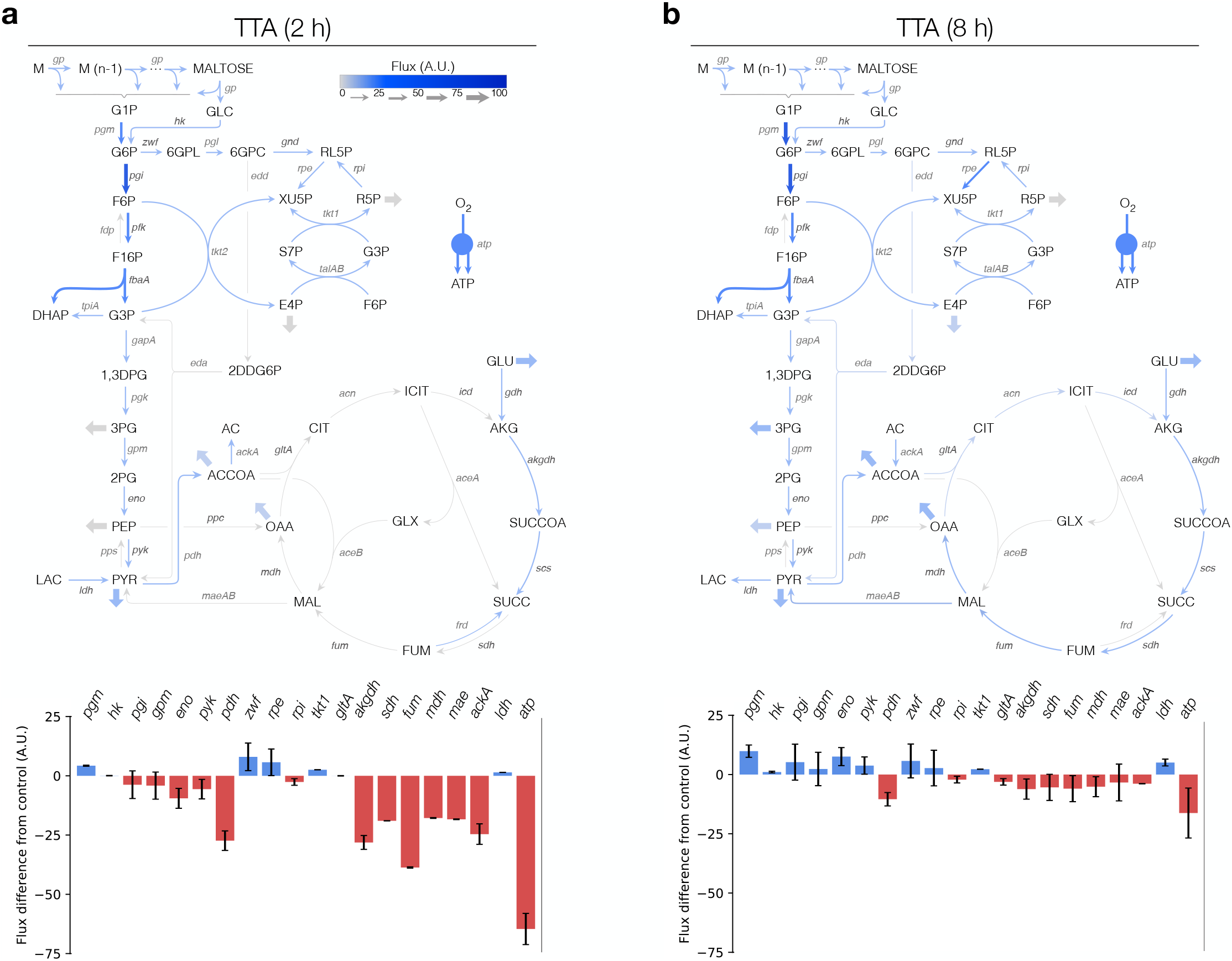
Mean flux distribution across an ensemble (N=100) for TTA. Fluxes were determined by integrating kinetic parameters with enzyme levels and constraining to measurements of metabolites and enzyme activity levels where data was available. (a) Flux distribution at 2 hours of CFPS reaction. Flux difference from control shown for key reactions at 2 hours of CFPS reaction. (b) Flux distribution at 8 hours of CFPS reaction. Flux difference from control shown for key reactions at 8 hours of CFPS reaction. Fluxes were normalized to maltodextrin consumption at t=0 hours.

### Optimality of cell-free systems

Flux balance analysis assumes the optimality of the translation rate, even following the addition of DNP or TTA, which introduced a metabolic perturbation altering the wild-type behavior of the cell-free extract. However, unlike *in vivo* systems, cell-free systems contain no active gene expression mechanisms to react to perturbations, thus, making them potentially sensitive to perturbations that could otherwise be regulated [38]. To explore the question of the optimality of a cell-free system, we used the method of minimization of metabolic adjustment (MOMA) to compute the optimal metabolic adjustment following the addition of the electron transport chain inhibitors DNP and TTA (Fig. 6). MOMA, developed by Church and coworkers [39], minimizes the distance between a presumably optimal nominal flux distribution and the flux distribution in the presence of a perturbation, in this case, treatment with DNP or TTA. For both inhibitors and the nominal case, an ensemble of N = 100 flux distributions was generated, given experimental noise and uncertainty in model parameters. The ensemble of flux balance solutions for the nominal treatment case (no inhibitors) was provided as an input to the MOMA calculation.

**Fig. 6:**
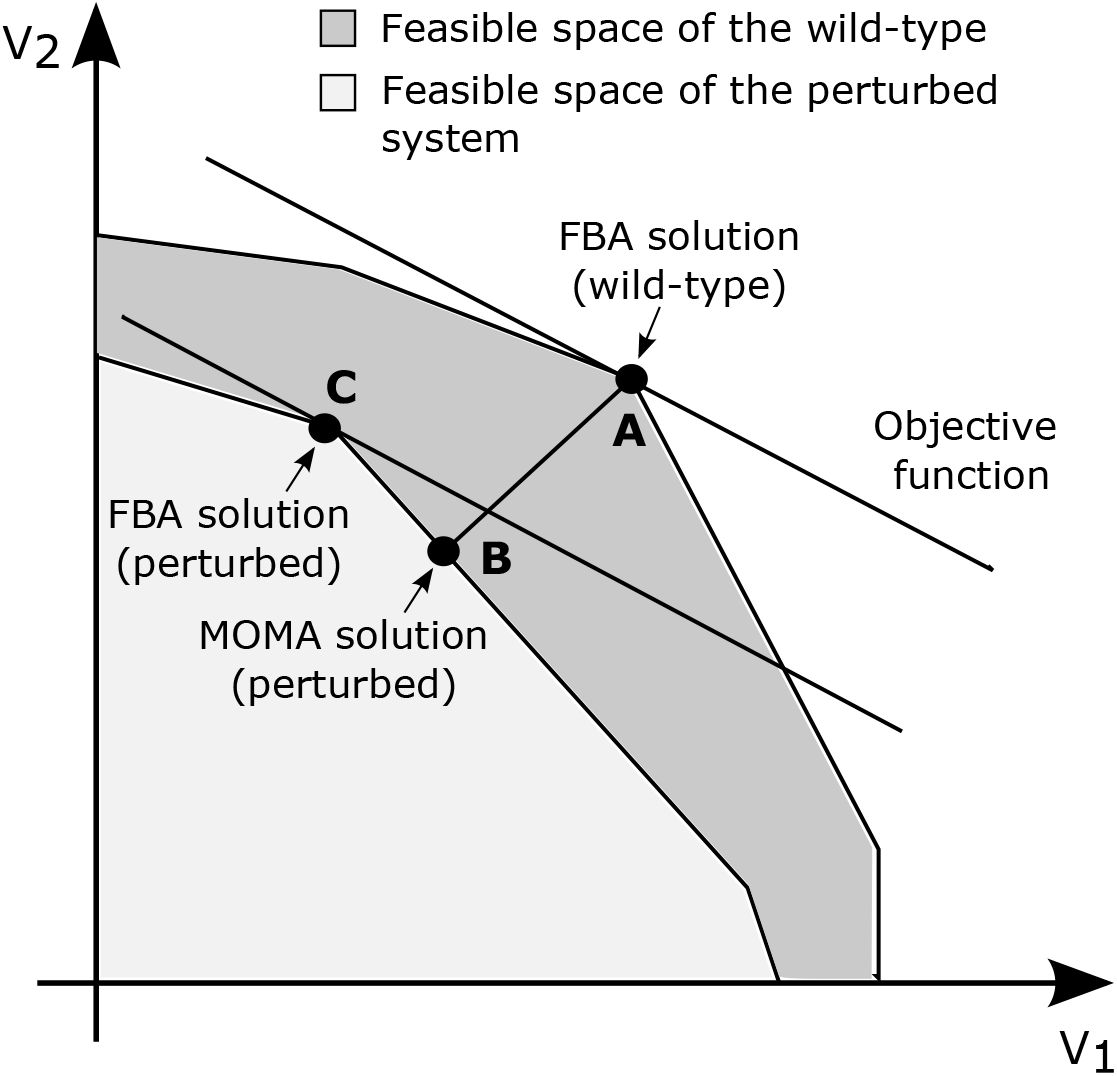
Schematic 2D representation of the feasible space for the wild-type (light grey polygon) and perturbed networks (dark grey polygon). The coordinates denote two arbitrary representative fluxes. Point **A** is the optimal FBA prediction for the wild type (or control case) and point **B** is the optimal FBA prediction for the perturbed system (with DNP/ TTA treatment). Point **C** is the alternative MOMA solution calculated through quadratic programming.

MOMA captured experimental mRNA and protein concentration measurements following DNP or TTA treatment, with only minor differences between the FBA and MOMA predictions (Fig. 7). In the case of DNP treatment, MOMA predicted a gradual increase in GFP protein accumulation between the 6- and 16-hour time points. On the other hand, MOMA failed to capture the mRNA concentration at the 16-hour time point following DNP treatment. Since MOMA minimizes the squared differences between the wild-type flux distribution and the unknown flux distribution, likely, the inconsistency between FBA simulations and experimental mRNA levels for the no-treatment case at the 16-hour time point affected MOMA predictions. In the case of TTA, the mean of the ensemble generated by MOMA marginally under-predicted experimental GFP protein levels. However, as the CFPS reaction progressed, the experimental data points fell within the predicted confidence interval. In summary, MOMA predicted GFP mRNA and protein production to occur under optimal conditions when the myTXTL cell-free system was incubated with DNP and TTA.

**Fig. 7:**
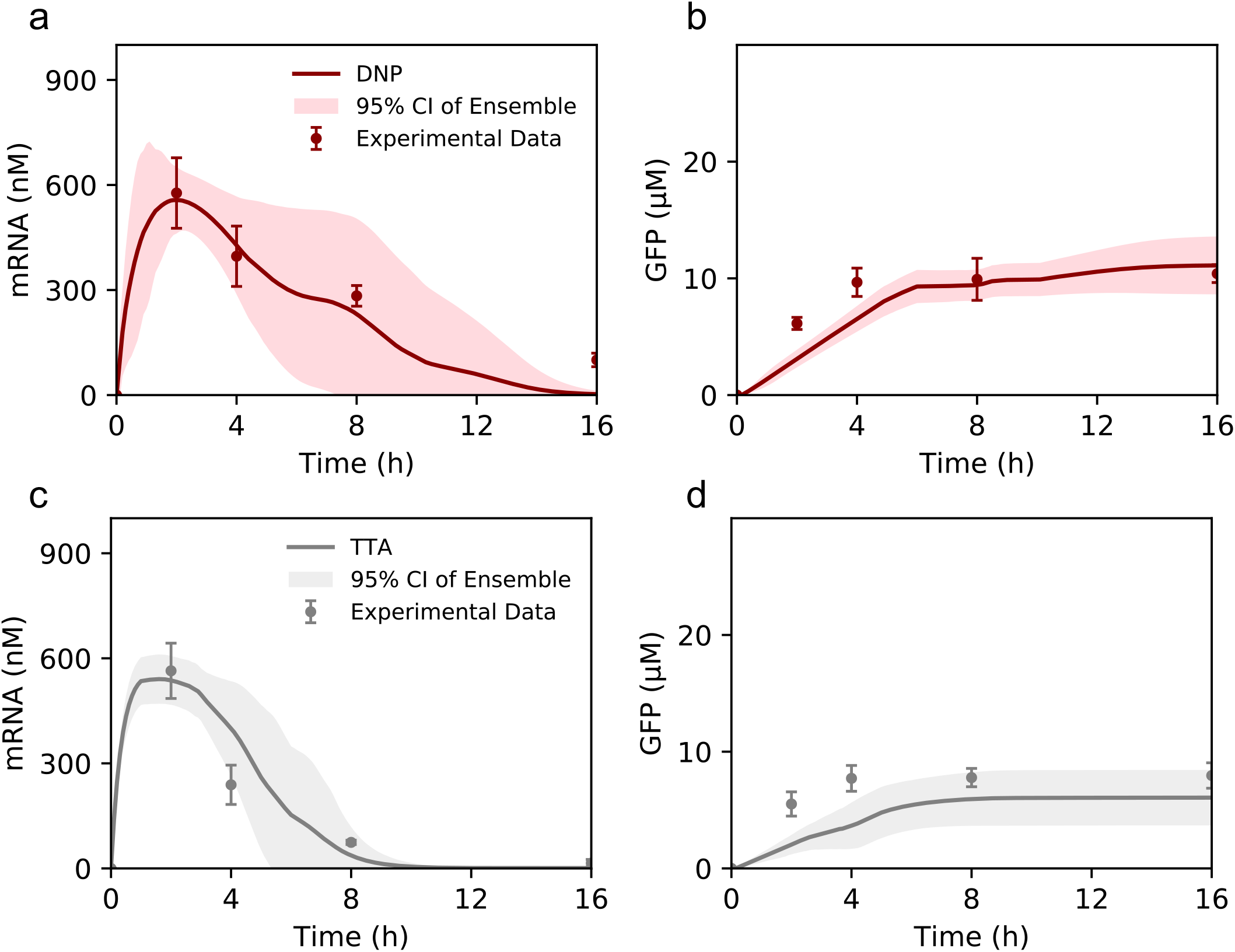
GFP production predicted using MOMA. (a) mRNA and (b) GFP protein predictions in the presence of DNP (red). (c) mRNA and (d) GFP protein predictions in the presence of TTA (grey). The shaded region denotes the 95% CI over an ensemble of N = 100 sets, the solid lines denote the mean of the ensemble, and points denote experimental measurements. Error bars denote the standard deviation of experimental measurements.

We compared the energy efficiency and carbon yield to understand the differences between the MOMA and FBA predicted flux distribution. MOMA predicted lower energy efficiency for the treated groups for transcription and translation compared to FBA (Fig. 8A-B). This is consistent with the mathematical requirement that FBA maximizes the translation rate for CFPS, therefore estimating higher energy utilization for GFP production. MOMA also predicted higher energy utilization for anaplerosis and amino acid biosynthesis than FBA for both the DNP and TTA cases. Otherwise, there were no significant differences between the FBA and MOMA solutions for the DNP case. However, for the TTA case, MOMA predicted almost 55% energy wasted toward degradation, much higher than the FBA prediction of 16%. Additionally, in the TTA case, MOMA predicted a much lower energy utilization in glycolysis compared to FBA (23% and 56% for MOMA and FBA, respectively). Thus, the simultaneous increase in energy utilization by amino acid biosynthesis and degradation pathways predicted by MOMA shows a more significant inefficiency in the distribution of energy resources across the metabolic network in the presence of TTA.

**Fig. 8:**
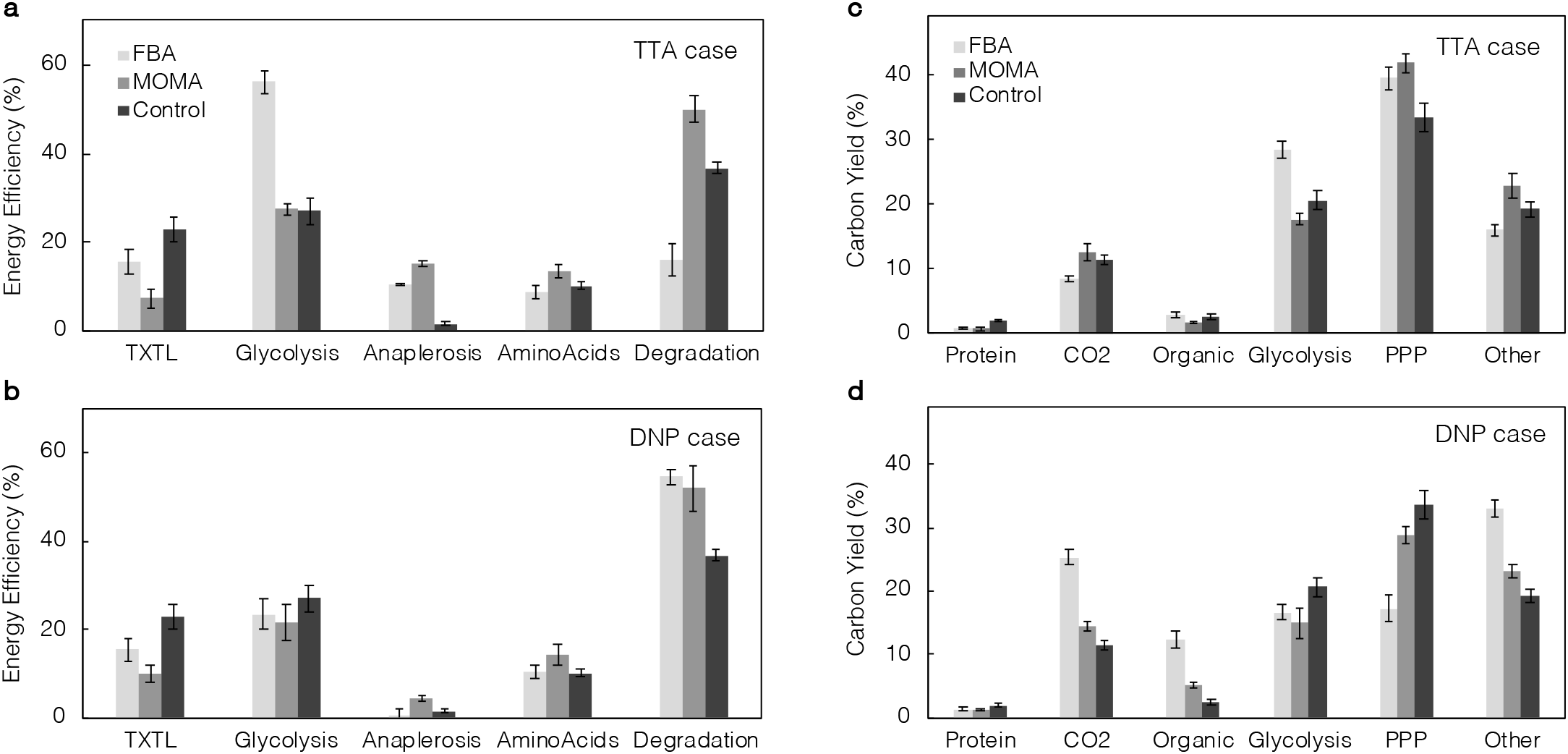
Performance metrics calculated from flux distributions predicted by FBA and MOMA. (a) Mean carbon yield across an ensemble (N=100) estimated for TTA and (b) DNP. CFPS carbon yield was calculated for the duration of the reaction as a ratio of the concentration of carbon produced to the total concentration of carbon consumed; PPP denotes the Pentose Phosphate Pathway and “Other” includes purine, pyrimidine, and chorismate metabolism. (c) Mean energy efficiency across an ensemble (N=100) estimated from FBA and MOMA simulations for TTA and (d) DNP. Energy efficiency was calculated as a ratio for the entire duration of the reaction in terms of nucleotide triphosphates utilized for the corresponding category to ATP generation.

Finally, we observed differences in the ultimate fate of carbon predicted by the FBA and MOMA objective functions (Fig. 8C-D). For DNP or TTA treatment, MOMA predicted significant carbon flux went toward the pentose phosphate pathway, followed by purine, pyrimidine, and chorismate metabolism (“other” category), glycolysis, and carbon dioxide. In addition, the MOMA prediction of carbon yield for GFP protein synthesis was marginally lower than FBA. Further, MOMA predicted reduced organic acid synthesis compared to FBA. This indicates that the cell-free extract in the MOMA simulations attempted to consume metabolites at wild-type levels and partially secreted the difference in the form of organic acids when they could not be consumed entirely. MOMA predicted a carbon flux distribution closer to the wild-type model compared to FBA for the treated groups. In summary, MOMA predicted GFP mRNA and protein production at optimal conditions with specific differences in the overall flux distribution.

## Discussion

The metabolism active during cell-free protein synthesis is not limited to transcription and translation processes. Jewett and coworkers have shown that central carbon metabolism is activated with inverted membrane vesicles carrying out oxidative phosphorylation in the Cytomim system, an *E. coli* based extract [15]. Thus, CFPS systems are often more complex than previously believed, with many interacting components that could be potentially exploited to optimize performance. In this study, we used a constraint-based modeling framework, in combination with a variety of metabolic measurements, to analyze the activity of metabolic pathways and biochemical reactions that were operating during the expression of a model protein in a commercially available *E. coli* cell-free extract.

This study generated and analyzed a comprehensive time-resolved data set for cellfree protein synthesis in the myTXTL system. A key advantage of a cell-free system is eliminating the cell membrane, which allows for direct access to metabolites and potentially precise control of metabolism. Toward this advantage, we presented a robust time-course quantification of metabolite concentration, the concentration of the 20 amino acids, and the absolute levels of mRNA and protein in the myTXLT cell-free system. This comprehensive dataset, which significantly extends previous studies [14, 15, 23], along with kinetic parameter values taken from BRENDA [34] and enzyme abundance estimated by Garenne and coworkers [36] (or assay reported in this study), allowed us to construct a dynamic constraint-based simulation of CFPS metabolism. The constraint-based model was validated by predicting mRNA and protein production in nominal and the DNP/TTA inhibitor treatment groups. DNP incubation has been reported to result in overconsumption of oxygen and a higher rate of metabolism while inhibiting energy-requiring processes [40, 41]. DNP has also been shown to disrupt oxidative phosphorylation in the Cytomim system [15], and in zebrafish, [42]. The model predicted a high rate of metabolism leading to acetate accumulation, agreeing with the literature findings. Further, the high carbon yield of 25% in carbon dioxide suggests higher activity metabolism and anaplerosis. TTA blocks the respiratory chain at Complex II by binding to the quinone reduction site and inhibiting the transfer of electrons from FE-S center S-3 of succinate dehydrogenase to oxidized ubiquinone [43, 44]. Jewett and coworkers also used TTA to assess whether oxidative phosphorylation was active in the Cytomim system. In our study, TTA incubation resulted in a lower protein titer of about 50% when compared to the control. The model predicted reduced activity in succinate dehydrogenase, and oxidative phosphorylation activity was decreased by 51% when compared to the control. Previously, the myTXTL system has been reported to rely on glycolysis and recycle inorganic phosphates [19, 45]; however, our study suggests that the *E. coli* based extract also has active oxidative phosphorylation with glutamate powering the TCA cycle.

Constraint-based modeling is a quantitative tool to assess and ultimately design the performance of a cell-free system. Previously, we reported a theoretical optimal energy efficiency of approximately 80% for transcription and translation [22]. However, the myTXTL system had an energy efficiency of 23% for the control and 15% for the inhibitors. In addition, the carbon yield for GFP was only 2%. Thus, CFPS has more than enough carbon and energy requirements but is not being effectively used. The flux distribution suggests that despite having oxidative phosphorylation, anaerobic processes are still active in cell-free extracts, as seen with the high accumulation of acetate and flux in anaplerotic reactions. Where *in vivo* systems can respond to different environmental conditions and activate different metabolic pathways, cell-free extracts no longer have the ability for enzymatic regulation. Thus, some of the enzymes present in CFPS may lead to inefficiency and low carbon yield. For example, Bujara and coworkers successfully increased the yield of dihydroxyacetone phosphate (DHAP) from glucose in CFPS [46]. The source strain used for cell-free extract preparation had a gene knockout of triosephosphate isomerase (*tpiA*) which resulted in the higher accumulation of DHAP. Such strategies have been used for decades in *in vivo* systems [47] and are only beginning to be used in CFPS [17, 48, 49]. In addition, most of the energy is wasted on nucleotide triphosphate degradation. Underwood and coworkers showed that increasing ribosome abundance did not significantly increase protein yields or rates; however, adding elongation factors increased protein synthesis rates by 27% [50], suggesting that the energy utilization could be optimized by addressing the rate-limiting step of protein production, namely translation [22, 51]. Subsequently, the carbon substrates that power CFPS could be minimized to increase the energy efficiency and carbon yield by lowering the total ATP produced since the majority is degraded.

The consistency of the constraint-based prediction of the mRNA and protein abundance with experimental measurements suggests that maximization of the translation rate is a potential objective function for genome-scale reconstructions of *E. coli* cell-free protein synthesis. Vilkhovoy et al. used the same objective in a different cell-free system and showed similar results [22]. However, the biological relevance of the selection of systemic optimality has always been questioned, especially for a perturbed system. Toward this question, we used the minimization of metabolic adjustment (MOMA) method to test the hypothesis of minimal re-adjustment of the wild-type cell-free flux distribution when challenged with a metabolic perturbation, namely treatment with electron transport chain inhibitors. MOMA predicted the overall mRNA and protein abundance in the presence of the electron transport chain inhibitors; however, there were differences between the FBA and MOMA predicted flux distributions. For example, analysis of energy efficiency under the MOMA objective showed an increase in the wastage of energy resources in the treatment groups through unnecessary side reactions. This was also reflected by decreased energy utilization for transcription and translation. These differences in flux distribution are potentially driven by the MOMA objective to match wild-type flux distribution; however, they could suggest the possibility of alternative optimal solutions despite the incorporation of experimental constraints. This is especially interesting because constraint-based models are under-determined. Thus, we expect different flux distributions satisfying the same conditions with similar CFPS productivity. Therefore, to ultimately determine the metabolic flux distribution occurring in CFPS, we could feed the system labeled substrates to resolve branch splitting estimations; ^13^C labeling techniques are well established for *in vivo* processes [52], however, the application of these techniques to cell-free systems remains an open area of research.

## Materials and Methods

### Cell-free protein synthesis and oxidative phosphorylation inhibitors

Cell-free protein synthesis reactions were carried out with the myTXTL system (Arbor Biosciences, Ann Arbor, MI) in 1.5 mL Eppendorf tubes at 29 °C. Plasmid P70a-GFP (Arbor Biosciences) was used as the DNA template for green flourescent protein (GFP) expression. The template plasmid was amplified in *E. coli* KL740 cI857+ (*E. coli* Genetic Stock Center, No. 14222). The plasmid was isolated and purified using a Plasmid Mini Kit (Qiagen, Valencia, CA). Each cell-free reaction was supplemented with a final concentration of 5 nM P70a-GFP. Each cell-free reaction had a total volume of 14 *μ*L with 9 *μ*L myTXTL master mix, 1.5 *μ*L P70a-GFP, and 3.5 *μ*L water (control), 3.5 *μ*L 2-4-dinitrophenol (DNP, 2.5 mM final concentration in CFPS), or 3.5 *μ*L thenoyltrifluoroacetone (TTA, 1 mM final concentration in CFPS). DNP was solubilized in water to prepare a 10 mM stock solution. TTA was solubilized in methanol to prepare a 100 mM stock solution, which was further diluted with water to 4 mM before being added to the CFPS reaction. Negative controls performed with methanol demonstrated that these solvents did not affect protein synthesis at concentrations used in this study. Separate CFPS samples were carried out in triplicate for each time point in order to ensure constant volume throughout the duration of the reaction.

### Absolute metabolite quantification

To quantify central carbon metabolites and amino acids, reaction samples were quenched with 100% ice-cold ethanol in a 1:1 volumetric ratio. Ethanol precipitated samples were centrifuged at 12,000 g for 15 minutes at 4 °C. The supernatant was collected and stored at -80 °C. Metabolites involved in glycolysis, pentose phosphate pathway, TCA cycle, and energy metabolism were quantified by liquid chromatography-mass spectrometry (LC-MS) using an isotope ratio based approach. Samples were tagged with 12C aniline, while internal standards were tagged with 13C aniline as described previously [53]. Briefly, 6 *μ*L of the supernatant was added to 44 *μ*L of water, followed by 5 *μ*L of 200 mg/mL EDC (N-(3-dimethylaminopropyl)-N-ethylcarbodiimide hydrochloride) and 5 *μ*L of 12C 6 M Aniline (pH 4.5). EDC was solubilized in water. The aniline solution was prepared by combining 550 *μ*L of 10.9 M aniline with 337.5 *μ*L water and 112.5 *μ*L of 12 M hydrochloric acid in an Eppendorf tube and vortexed well. The mixture was gently vortexed at room temperature for 2 hours. In order to stabilize the metabolites, 1.5 *μ*L of TEA (triethylamine) was added. The mixture was centrifuged at 13,500 g for 3 minutes and 25 *μ*L of the supernatant was transferred to a LC-MS vial. The sample mixture was mixed with 25 *μ*L of a standard stock solution containing 35 metabolites at 80 *μ*M tagged with 13C aniline. The standard stock solution was tagged with aniline following the same procedure as the sample, except with 13C aniline. The 35 standard metabolites are listed in the metabolite dataset and exclude acetic acid, NAD, NADP, FAD, acetyl-CoA, glycerol 3-phosphate, and maltose.

Acetic acid was tagged with aniline and quantified with a standard curve method. NAD, NADP, FAD, acetyl-CoA and glycerol 3-phosphate were not tagged with aniline, and were quantified by a standard curve method. Samples and standards were injected at 5 *μ*L onto a Waters Acquity BEH C18 (1.7 *μ*m, 2.1 mm x 150 mm) column. The LC-MS system consisted of a Waters Acquity Quaternary system, an Acquity Sample Manager, and an Acquity QDa detector (Waters Corp, Medford, MA). The system was controlled by Empower 3 software (Waters). The autosampler was set at 10 °C. Separation was carried out at a flow rate of 0.3 mL/min. The elution started with 95 % mobile phase A (5 mM tributylamine (TBA) in HPLC-grade water adjusted to pH 4.75 with glacial acetic acid) and 5 % mobile phase B (5 mM TBA in acetonitrile), raised to 70 % B in 10 minutes, raised to 100 % B in 2 minutes and held at 100 % B for 3 minutes. Return to initial conditions (95 % A, 5 % B) over 1 minute and hold for 9 minutes to re-equilibrate the column. The column was pre-conditioned with the specified gradient protocol 3 times prior to any injection onto the column. The MS chromatograms were acquired in negative ion mode with a probe temperature of 520 °C, negative capillary voltage of -0.8 kV, and positive capillary voltage of 0.8 kV.

### Amino acid analysis

Amino acids were analyzed using a Waters AccQ-Tag Ultra Amino Acid Analysis Kit (Waters). The ethanol-precipitated CFPS samples were derivatized and tagged by combining 4 *μ*L of the sample with 6 *μ*L of water, 70 *μ*L borate buffer solution (Waters), and 10 *μ*L reagent (Waters). The solution was then kept in a water bath at 55 ° C for 10 minutes. One microliter was then injected onto an Acquity BEH C18 column (1.7 *μ*m, 2.1 mm x 100 mm). The elution gradient and flow rate were set according to the manufacturer’s recommendations. Amino acids were detected by an Acquity TUV detector (Waters) at 260 nm. Amino acids were identified by known retention times of standards. Concentrations were determined by comparison with calibration standard curves with the exception of glutamate. Glutamate was outside the linearity of the calibration standard curves and not quantified with the Waters AccQ-Tag Ultra Amino Acid Analysis Kit.

### Glutamate and maltose assays

Glutamate and maltose concentrations were determined using enzymatic colorimetric assays purchased from Sigma-Aldrich (St. Louis, MO) according to the manufacturer’s instructions. The readings were performed with a multimode plate reader Varioskan Lux (ThermoFisher, Waltham, MA) using 96-well plates.

### Protein quantification

GFP concentrations were determined by fluorescence measurements with comparison to a standard curve. Two microliters of each CFPS reaction were diluted with 33 *μ*L of phosphate-buffered saline and analyzed in triplicate on a black 384-well plate with a multimode plate reader Varioskan Lux (ThermoFisher) at 488 nm excitation and 535 nm emission.

### Enzyme activity assays

Enzyme activity for 6-phsophogluconate dehydrogenase (*gnd*) and phosphoenolpyruvate carboxylase (*ppc*) were determined using colorimetric based assays purchased from BioVision (Milpitas, CA) according to the manufacturer’s instructions. All remaining enzyme activity levels were determined using colorimetric and fluorescence based assays purchased from Sigma-Aldrich (St. Louis, MO) according to the manufacturer’s instructions. The readings were performed in kinetic (loop) mode with a multimode plate reader Varioskan Lux (ThermoFisher) using clear 96-well plates for colorimetric assays and black 96-well plates for fluorescence based assays.

### Absolute quantification of mRNA

Absolute levels of messenger RNA (mRNA) were quantified using quantitative real-time RT-PCR with comparison to a standard curve. A standard of GFP mRNA was prepared by conducting a CFPS reaction with 5 nM P70a-GFP plasmid for 2 hours at 29 °C. The reaction was applied to a PureLink RNA Mini Kit with an on-column PureLink DNase Treatment (ThermoFisher) according to the manufacturer’s instructions. The total RNA was eluted with Invitrogen UltraPure DNase/RNase-free water (ThermoFisher). The total RNA was then applied to MICROBExpress Bacterial mRNA Enrichment Kit (ThermoFisher) followed by MEGAclear Transcription Clean-up Kit (ThermoFisher) according to the manufacturer’s instructions. The purified mRNA was eluted with UltraPure water. The mRNA concentration was determined with a Qubit Fluorometer using a Qubit RNA HS Assay Kit (ThermoFisher).

To quantify mRNA levels in CFPS samples from the experiment, 1 *μ*L from the CFPS reaction was applied to the PureLink RNA Mini Kit with an on-column PureLink DNase Treatment according to the manufacturer’s instructions. The total RNA was eluted with 50 *μ*L of UltraPure water. The total RNA sample was diluted 100-fold, and 2 *μ*L of the diluted sample was loaded for each RT-PCR reaction. The quantitative real-time RT-PCR reaction was carried out on an Applied Biosystems QuantStudio 3 with a Taqman RNA-to-C_t_ 1-Step Kit using GFP Taqman assay (Mr04329676 mr) on a 96-well plate in triplicate according to the manufacturer’s instructions (Applied Biosystems, Life Technologies Corporation, Foster City, CA). Sample mRNA concentrations were determined by comparison to the calibration standard curve. The standard curve was generated with the purified mRNA of GFP ranging from 10−4 to 1 ng. The standard curve had a linearity coefficient of 0.994 and efficiency of 104 %.

### Flux balance analysis

The dynamic sequence specific flux balance analysis problem was formulated as a linear program:

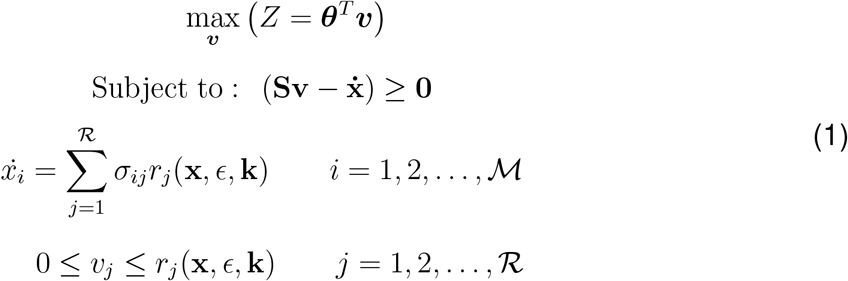

where **S** denotes the stoichiometric matrix (ℳ × ℛ) and *σ*_*ij*_ denotes the stoichiometric coefficient for species *i* in reaction *j*, **v** denotes the unknown flux vector (ℛ × 1), ***θ*** denotes the objective vector (ℛ × 1), and *r*_*j*_(**x**, *ϵ*, **k**) denotes the rate of reaction *j*. For all metabolic reactions except for the transcription/translation processes and maltodextrin consumption, reaction *j* was modeled as the product of the turnover rate *k*_*j*_ and enzyme abundance *ϵ*_*j*_ or known as the maximum velocity of the reaction *V*_*max*_ (mM/h). The transcription/translation and maltodextrin consumption reactions were modeled following saturation kinetics. The turnover rate for each reaction was identified from BRENDA [34] or taken from Adadi and coworkers [35]. The enzyme abundance was identified for 104 reactions from Garenne and coworkers [36]. Garenne and coworkers reported the counts of the enzymes identified in their LC-MS analysis, where we calibrated the counts of sigma 70 and RNA polymerase to the concentration values [19] to create a calibration curve. The enzyme abundance was calculated using this calibration curve and all remaining enzymes not identified in Garenne and coworkers were set to the median value of 50 nM. The enzyme abundance was validated for a subset of 15 enzymes from our enzyme activity assays where we calculated the expected enzyme abundance in the cell-free reaction by:

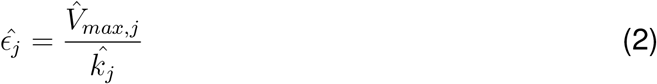

where 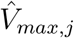 is the maximum velocity for reaction *j* from the enzyme activity assay and 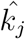 is the corresponding turnover number for enzyme *j*.

### Minimization of metabolic adjustment (MOMA)

The MOMA formulation used the same constraints as flux balance analysis but relaxed the assumption of optimality of a specific biological objective. MOMA was formulated as a quadratic program [39]:

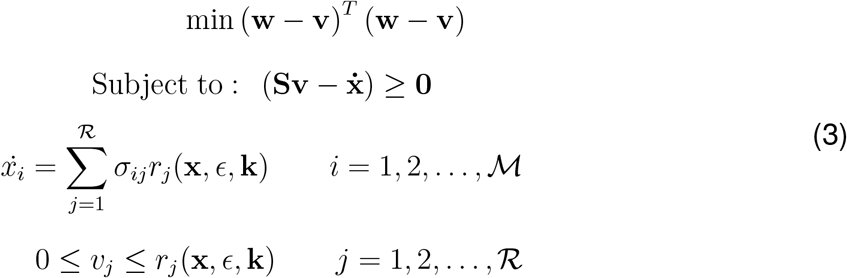

where **w** denotes the known wild type flux vector (control case) and **v** denotes the unknown flux vector of the perturbed system (with TTA or DNP treatment). The time-course flux distribution for the control case obtained from flux balance analysis was used as the wild-type flux distribution input to MOMA. The quadratic programming problem was set up using the Convex.jl [54] Julia package, which was solved using the Gurobi optimizer [55].

### Transcription and translation stoichiometric and kinetics

The transcription (TX) and translation (TL) reactions stoichiometry was modeled based on previous work [22, 28]. The transcription initiation rate was modeled as:

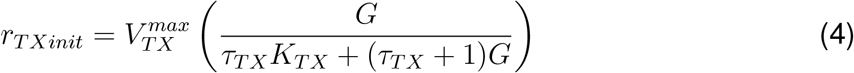

where *G* denotes the concentration of the DNA plasmid in the cell-free reaction, *K*_*TX*_ denotes a transcription saturation coefficient, and *τ*_*TX*_ denotes the transcription constant. The maximum transcription rate 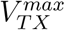 was formulated as:

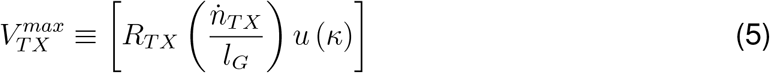

where *R*_*TX*_ denotes the RNA polymerase concentration, 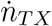 denotes the RNA polymerase elongation rate (nt/h), *l*_*G*_ denotes the gene length (nt). The term *u* (*κ*) (dimensionless, 0 ≤ *u* (*κ*) ≤ 1) is an effective model of promoter activity, where *κ* denotes promoter specific parameters. In this study, the promoter model was taken from Vilkhovoy and coworkers [22] for the P70a promoter. The transcription rate was modeled as:

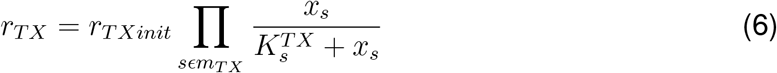

where *m*_*TX*_ denotes the set of reactants for transcription: ATP, CTP, GTP, and UTP, and 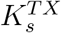 denotes the saturation constant for species *s*. The degradation of mRNA was modeled as a first order rate:

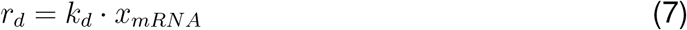

where *k*_*d*_ denotes the degradation rate constant. The translation initiation and translation rate was modeled as:

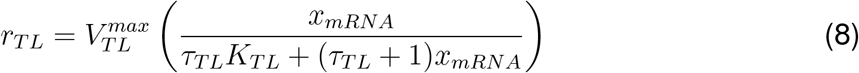

where *x*_*mRNA*_ denotes the concentration of the mRNA, *K*_*TL*_ denotes a translation saturation coefficient, and *τ*_*TL*_ denotes the translation constant. The maximum translation rate 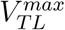 was formulated as:

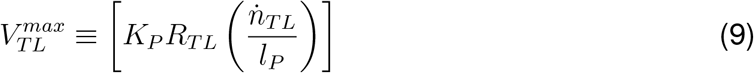

where *K*_*P*_ denotes the polysome amplification constant, *R*_*TL*_ denotes the ribosome concentration, 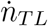 denotes the ribosome elongation rate (amino acids per hour), and *l*_*P*_ denotes the number of amino acids in the protein of interest. The abundance of each species *x* was modeled as:

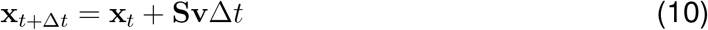

where *t* denotes the current time point and Δ*t* denotes the time step. Lastly, we imposed a user configurable bound ℬ_*i*_ on the maximum rate of change for metabolite *i* where data was available:

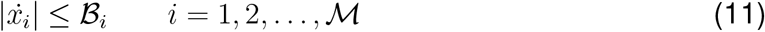

The bound ℬ_*i*_ was determined by fitting the time-course concentration data by a regression spline with the cubic SmoothingSplines package in Julia 1.1. The rate of change at time step *t* was determined by a forward difference approximation from *t* to *t* + Δ*t* from the regression spline. Metabolic fluxes were estimated at each time step using the GNU Linear Programming Kit (GLPK) v4.55 [56]. All parameters are listed in Table 1. In addition, flux bounds were set to the experimental value where data was available for the corresponding enzyme activity assays. The objective of the cell-free flux balance calculation was to maximize the rate of maltodextrin consumption, transcription initiation, transcription, mRNA degradation, translation initiation and translation, unless specified.

**Table 1:** Parameters for sequence specific flux balance analysis

| Description | Parameter | Value | Units | Reference |
| --- | --- | --- | --- | --- |
| RNA polymerase concentration | $R_{TL}$ | 60-75 | nM | [19] |
| Ribosome concentration | $R_{TX}$ | 2-2.3 | μM | [19] |
| Transcription elongation rate | $\dot{n}_{TX}$ | 15-25 | nt/s | [19] |
| Translation elongation rate | $\dot{n}_{TL}$ | 1-2 | aa/s/ribosome | [19, 50] |
| Transcription time constant | $\tau_{TX}$ | 0.021 - 0.05 | constant | calculated |
| Translation time constant | $\tau_{TL}$ | 0.063 - 0.126 | constant | calculated |
| Transcription saturation coefficient | $K_{TX}$ | 0.3 | μM | [57] |
| Translation saturation coefficient | $K_{TL}$ | 600.0 | μM | estimated |
| Polysome number | $K_P$ | 10 | ribosome number | estimated |
| mRNA degradation rate constant | $k_d^{mRNA}$ | 2.38 | $h^{-1}$ | [19] |
| Maltodextrin saturation constant | $K_m$ | 8.3 | mM | BRENDA |
| Transcription saturation constant | $K_s^{TX}$ | 0.03 | mM | estimated |
| Weight RNA polymerase binding alone P70a | $K_1$ | 0.014 | constant | estimated |
| Weight bound RNAP-$\sigma_{70}$ P70a | $K_2$ | 10 | constant | estimated |
| $\sigma_{70}$ concentration | $\sigma_{70}$ | 35 | nM | [19] |
| $\sigma_{70}$ dissociation constant | $K_D$ | 130 | nM | [58] |
| $\sigma_{70}$ hill coefficient | $n$ | 1 | constant | [58] |
| Gene concentration | $G_P$ | 5 | nM | experiment |
| ATP transcription coefficient | $ATP_{TX}$ | 208 | constant | calculated |
| CTP transcription coefficient | $CTP_{TX}$ | 157 | constant | calculated |
| GTP transcription coefficient | $GTP_{TX}$ | 195 | constant | calculated |
| UTP transcription coefficient | $UTP_{TX}$ | 157 | constant | calculated |
| ATP tRNA charging coefficient | $ATP_{TL}$ | 239 | constant | calculated |
| GTP translation coefficient | $GTP_{TL}$ | 478 | constant | calculated |
| Carbon number of GFP | $C_{GFP}$ | 1208 | constant | calculated |

### Quantification of uncertainty

Experimental factors taken from literature, for example macromolecular concentrations or elongation rates, are uncertain. To quantify the influence of this uncertainty on model performance, we randomly sampled the expected physiological ranges for these parameters as determined from literature. An ensemble of flux distributions was calculated for the three different cases we considered: control, DNP, and TTA. The flux ensemble was calculated by randomly sampling the rate of change for metabolites where data was available, randomly sampling enzyme abundance, and randomly sampling RNA polymerase levels, ribosome levels, and elongation rates in a physiological range determined from literature. The rate of change for metabolites was sampled between the calculated value from the regression spline to twice its value. The enzyme abundance was randomly sampled from the estimated value up to 1.5 times its value. P70 RNA polymerase levels were sampled between 60 and 75 nM, ribosome levels between 2.0 and 2.3 μM, the RNA polymerase elongation rate between 15 and 25 nt/s, and the ribosome elongation rate between 1.0 and 2.0 aa/s [19, 50]. We generated uniform random samples between an upper (*u*) and lower (*l*) parameter bound of the form:

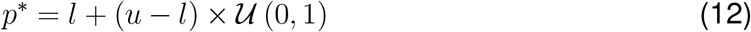

### Calculation of energy efficiency

Energy efficiency (ℰ) was calculated as the ratio of transcription and translation (weighted by the appropriate energy species coefficients) to ATP generation:

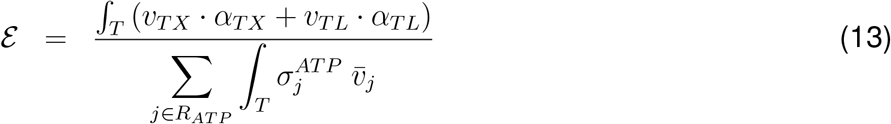

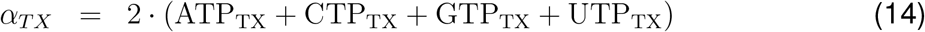

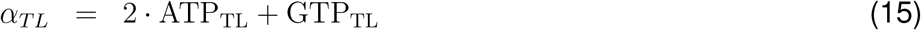

where *α*_*TX*_ denotes the energy cost of transcription, *α*_*TL*_ denotes the energy cost of translation, *R*_*ATP*_ denotes the set of ATP-producing reactions, 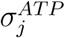 denotes the ATP coefficient for reaction *j*, and *T* denotes the time of the experiment. ATP_TX_, CTP_TX_, GTP_TX_, and UTP_TX_ denote the stoichiometric coefficients of each energy species for the transcription of the protein of interest, and ATP_TL_ and GTP_TL_ denote the stoichiometric coefficients of ATP and GTP for the translation of the protein of interest. During transcription and tRNA charging, triphosphate molecules are consumed with monophosphates as byproducts; this is the reason for the factors of 2 on ATP_TX_, CTP_TX_, GTP_TX_, UTP_TX_, and ATP_TL_.

### Calculation of carbon yield

The carbon yield (*Y*_*C*_) was calculated as the ratio of carbon produced as the protein divided by the carbon consumed as reactants:

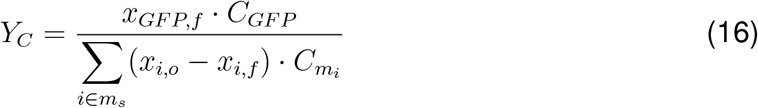

where *x*_*GFP,f*_ denotes the final concentration of GFP, *C*_*GFP*_ denotes carbon number of GFP, *m*_*s*_ denotes the set of species that were consumed, *x*_*i,o*_ denotes the initial concentration of species *i, x*_*i,f*_ denotes the final concentration of species *i*, and 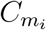 denotes the carbon number of species *i*.

## Author’s contributions

J.V directed the study. M.V., S.V. and A.A. developed the FBA model and designed and conducted the metabolomic and mRNA/protein quantification experiments. S.D. developed the MOMA model and analyzed the MOMA simulations.

## Acknowledgements

We gratefully acknowledge the suggestions from the anonymous reviewers to improve this manuscript.

## Data availability statement

Model code is available at: https://github.com/varnerlab/CFPS-FBA-MOMA-Code.git. The mRNA, protein, and metabolite measurements presented in this study are available in the data directory of the model repository in comma-separated value (CSV) and Microsoft Excel format.

## Supplementary Information

### Allosteric enzyme regulation mechanisms were active in CFPS

Allosteric regulation mechanisms were active in the myTXTL CFPS system (Fig. S1). The activity of enzymes throughout central carbon metabolism was measured for the control, DNP, and TTA groups at two and eight hours. Significant differences were observed between groups and the two-time points for several enzymes. However, for enzymes where allosteric regulation was not present, including *eno, mdh, gdh*, and *akgdh*, enzyme activity measurements were similar between groups and time points. Thus, the activity assays suggested that allosteric regulation was likely present in CFPS. For instance, the enzyme *icd* is allosterically inhibited by phosphoenolpyruvate (PEP). The control and TTA groups showed the enzyme activity increased from approximately 180 to 250 mM/h and 125 to 290 mM/h, respectively, from 2 to 8 hours. Concurrently, PEP abundance decreased for the control and TTA groups in the CFPS extract from 2 to 8 hours, which likely resulted in increased enzyme activity for *icd*.

### Network efficiency was altered in the presence of electron transport chain inhibitors

Energy efficiency was calculated as a ratio of nucleotide triphosphates utilized to ATP generation (see materials and methods). Energy efficiency was significantly higher for the control at 23 *±* 2.8% for transcription and translation, while the DNP and TTA group were 15 *±* 2.7% and 15 *±* 2.6%, respectively (Fig. S6). Despite having a higher energy efficiency than the treatment groups, 37 *±* 1% of the nucleotide triphosphates were wasted toward degradation (Fig S6A). Twenty-seven percent of energy was utilized in glycolysis, with 11% toward amino acid biosynthesis and 2% toward anaplerosis. In the DNP case, 50% of the energy was wasted toward degradation. This is due to DNP’s effect, which resulted in higher metabolism and the inhibition of all energy-requiring processes [41]. For TTA, the majority (51%) of the energy was spent on glycolysis. The flux distribution for TTA showed only an active upper glycolysis pathway and thus resulted in higher-than-normal energy utilization.

Carbon yield was calculated for the duration of the reaction as a ratio of the carbon produced to the total carbon consumed. Carbon yield for GFP production was similar for all treatment groups (Fig. S7). The control group had a carbon yield of 2 *±* 0.2%, whereas the DNP group had a carbon yield of 1.4 *±* 0.2% and TTA had a carbon yield of 1 *±*0.2 %. The low carbon yield for all groups showed that the myTXTL system was supplemented with more carbon than needed. For instance, the extract is supplied with 20-40 mM of maltodextrin and was measured to have a concentration of approximately 104 mM of glutamate. Meanwhile, the amount of GFP produced was in the range of 10-30 *μ*M. Thus, we investigated where the remaining carbon went. The control and TTA groups had very similar distributions of carbon in the network. Between both groups, 9-12% went toward carbon dioxide, 22-28% remained in glycolysis, 3% remained in the TCA cycle, 36-39% remained in the pentose phosphate pathway, 1-3% went toward amino acid biosynthesis, and 17-24% went toward purine, pyrimidine and chorismate metabolism (other category). Meanwhile, the carbon yield in the DNP group had a more uniform distribution throughout the network, with a notable difference in carbon dioxide (25%). The higher carbon dioxide output further supports the higher metabolism observed with DNP incubation [41].

**Fig. S1:**
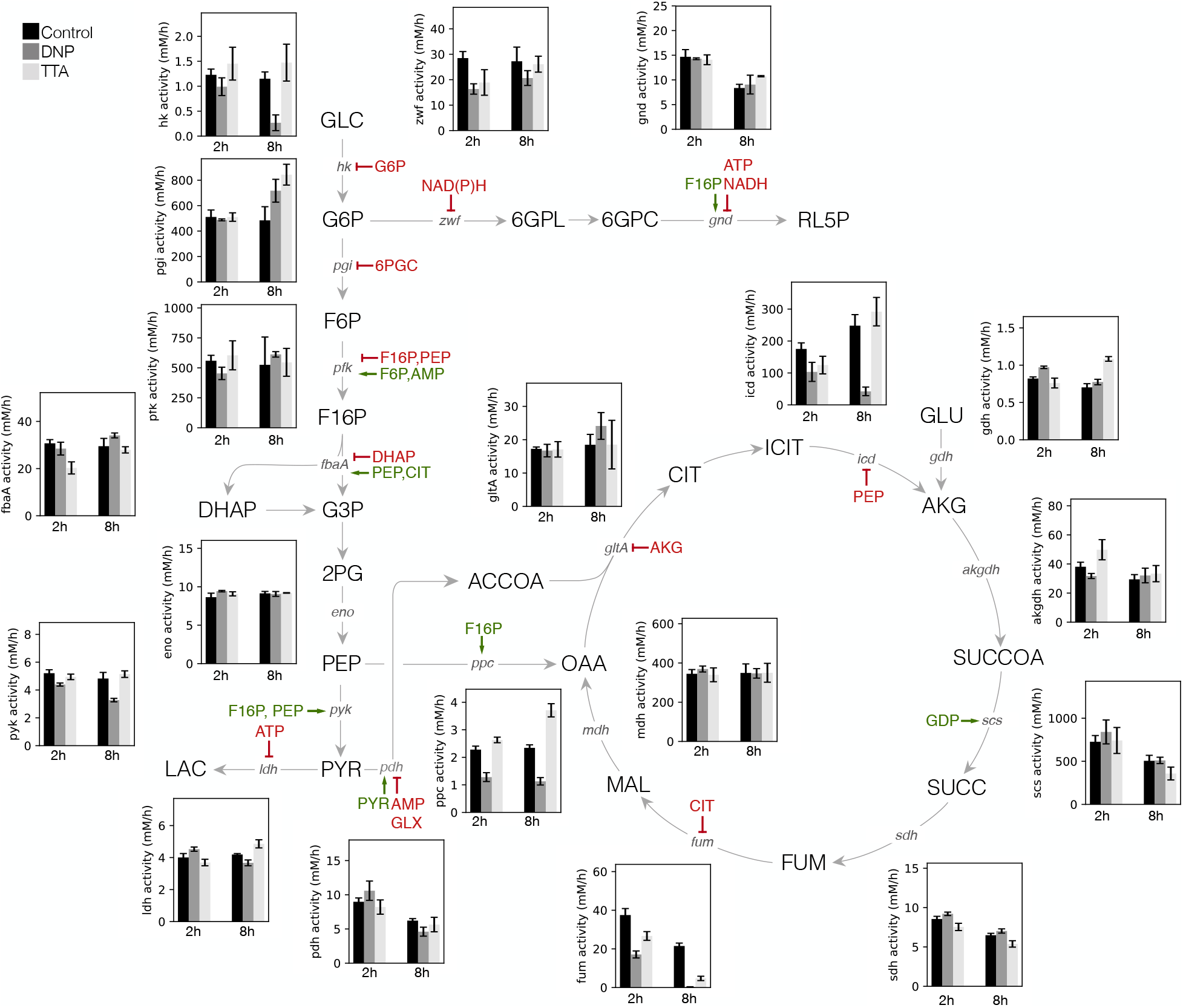
Enzyme activity measurements reveal allosteric regulation is present in CFPS. Enzyme activity assays at 2 and 8 hours of the CFPS reaction throughout the metabolic network for control (black), DNP (dark grey), and TTA (light grey).

**Fig. S2:**
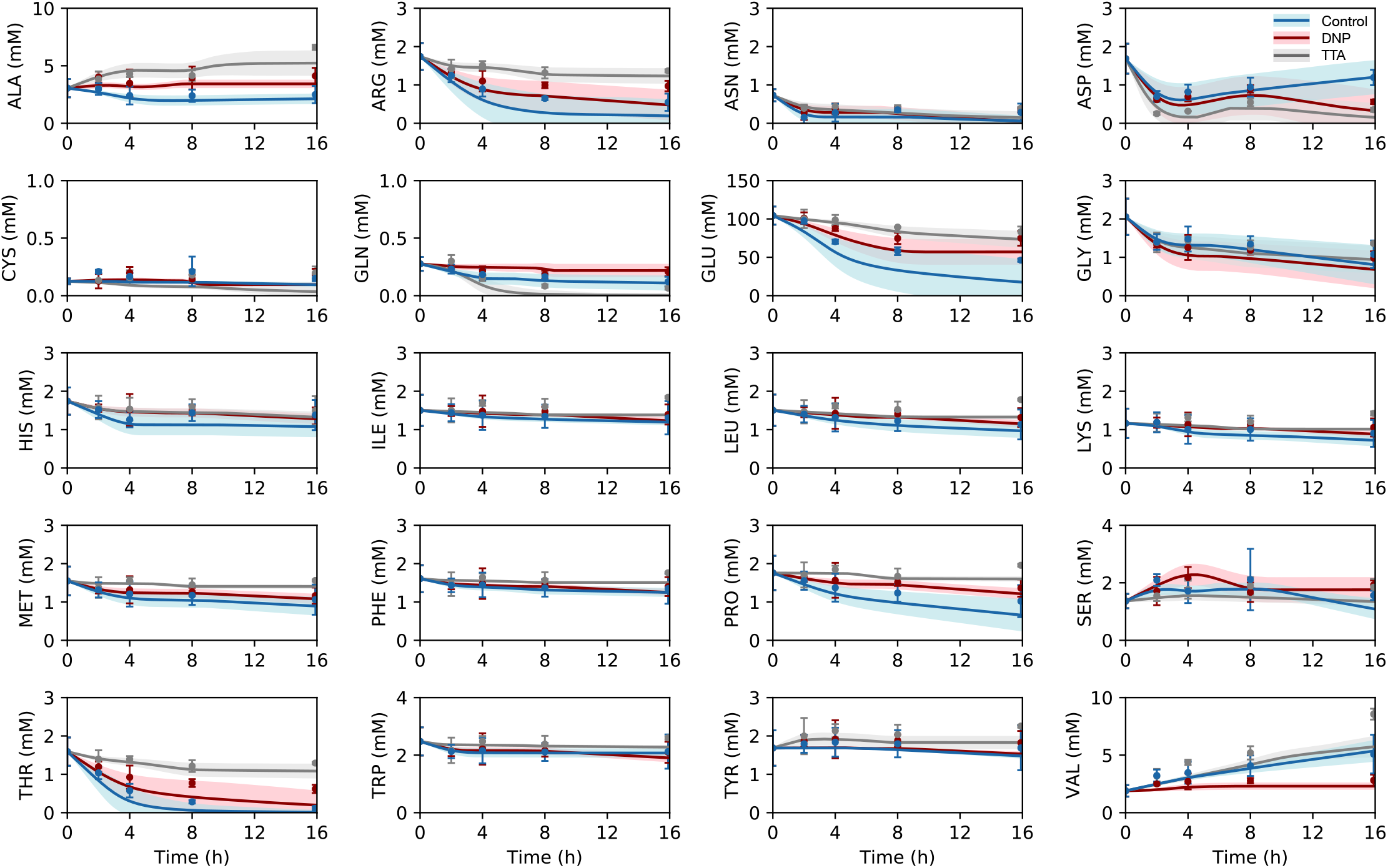
Time course of amino acid levels in CFPS for control (blue), DNP (red), and TTA (grey). Experimental amino acid fluxes constrained the mathematical model of CFPS. The solid line denotes the mean of the ensemble (N=100), the shaded region denotes the 95% confidence interval of the ensemble, the points denote experimental measurements, and error bars denote the standard deviation of experimental measurements.

**Fig. S3:**
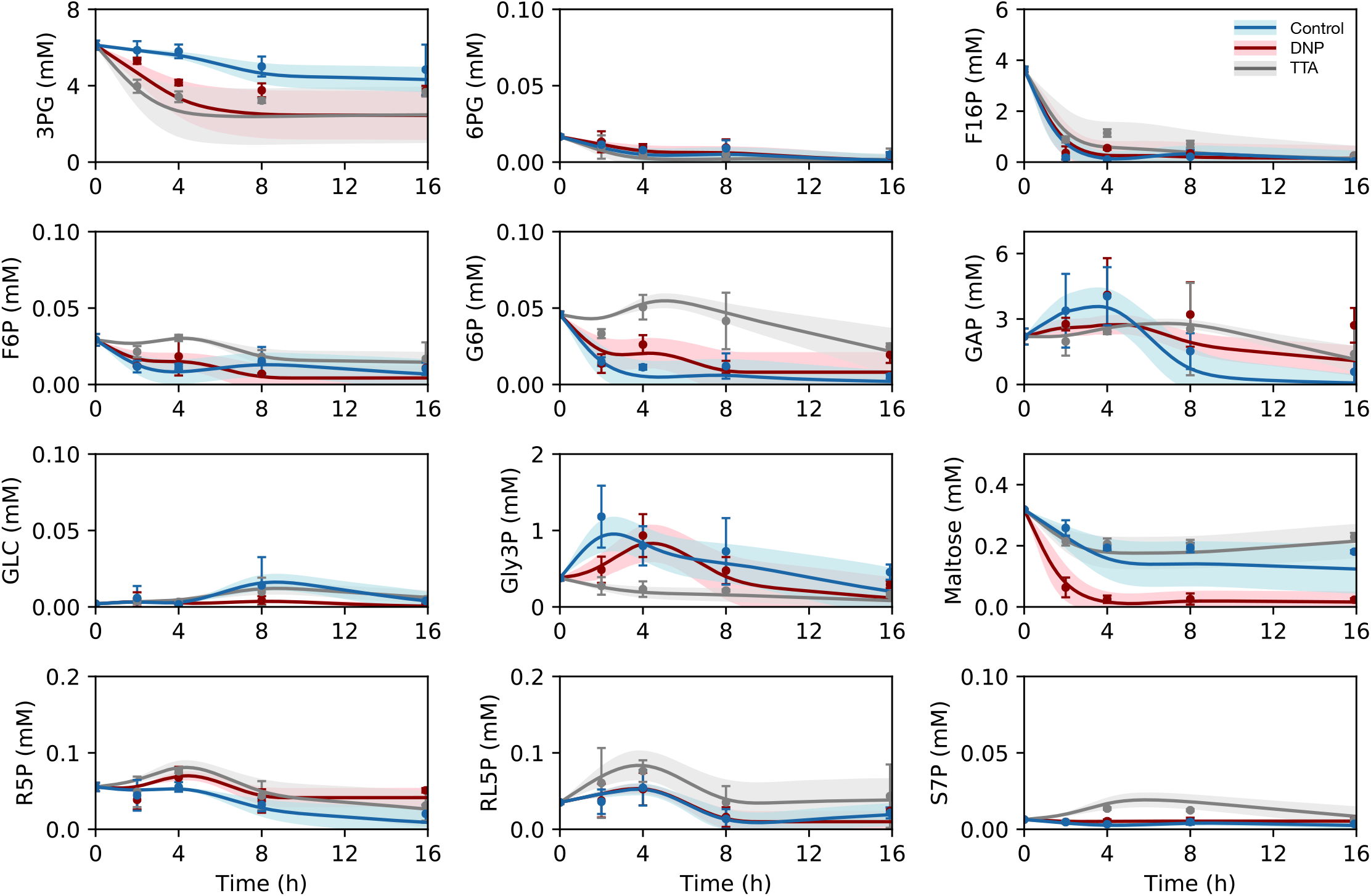
Time course of upper central carbon metabolite levels in CFPS for control (blue), DNP (red), and TTA (grey). DNP showed exhaustion of maltose revealing maltodextrin depletion and thus high carbon utilization. Experimental fluxes constrained the mathematical model of CFPS. The solid line denotes the mean of the ensemble (N=100), the shaded region denotes the 95% confidence interval of the ensemble, the points denote experimental measurements, and error bars denote the standard deviation of experimental measurements.

**Fig. S4:**
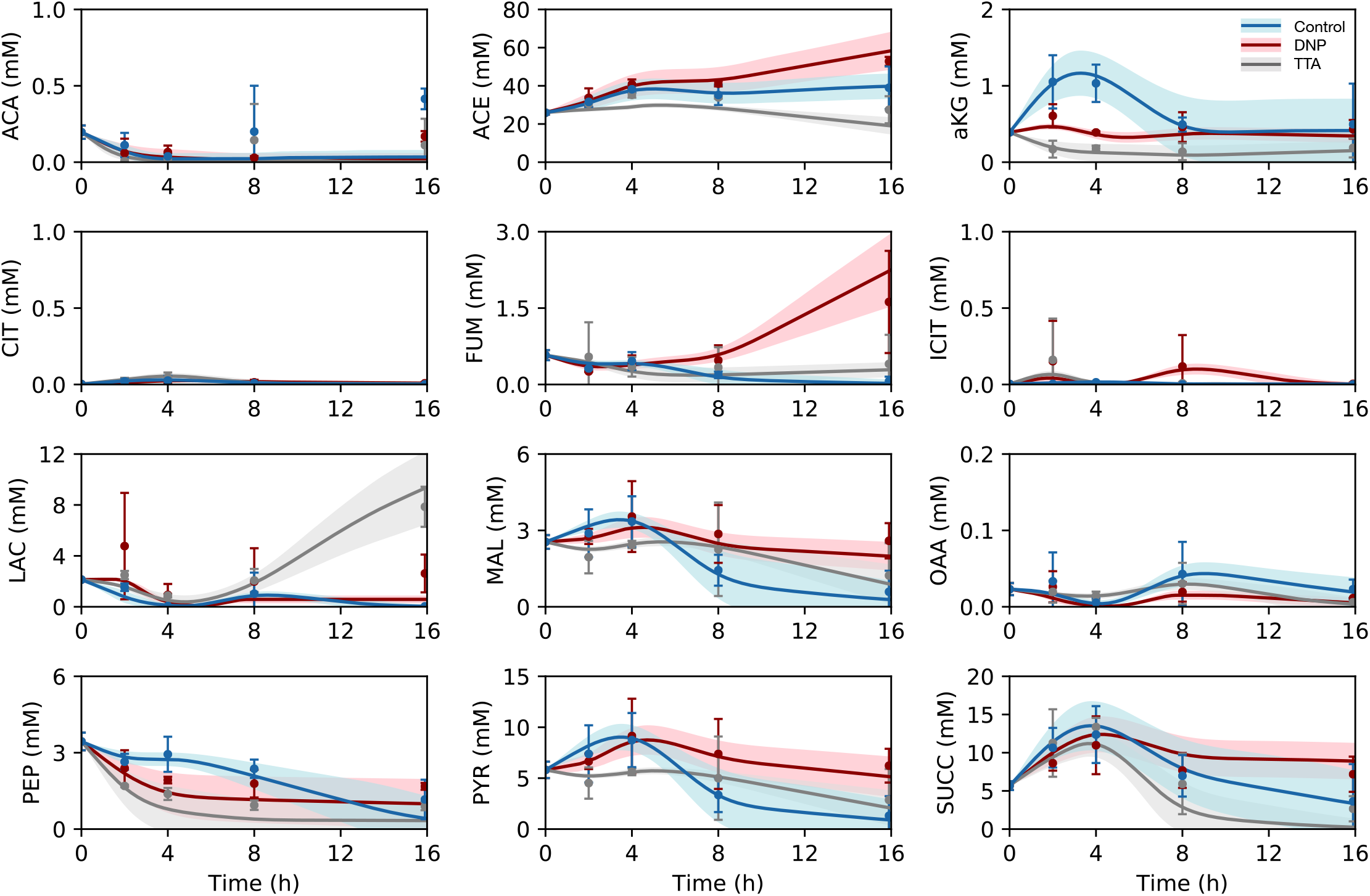
Time course of lower central carbon metabolite levels in CFPS for control (blue), DNP (red), and TTA (grey). DNP heavily relied on substrate level phosphorylation with high accumulation of acetate, whereas TTA had a high abundance of lactate. Experimental fluxes constrained the mathematical model of CFPS. The solid line denotes the mean of the ensemble (N=100), the shaded region denotes the 95% confidence interval of the ensemble, the points denote experimental measurements, and error bars denote the standard deviation of experimental measurements.

**Fig. S5:**
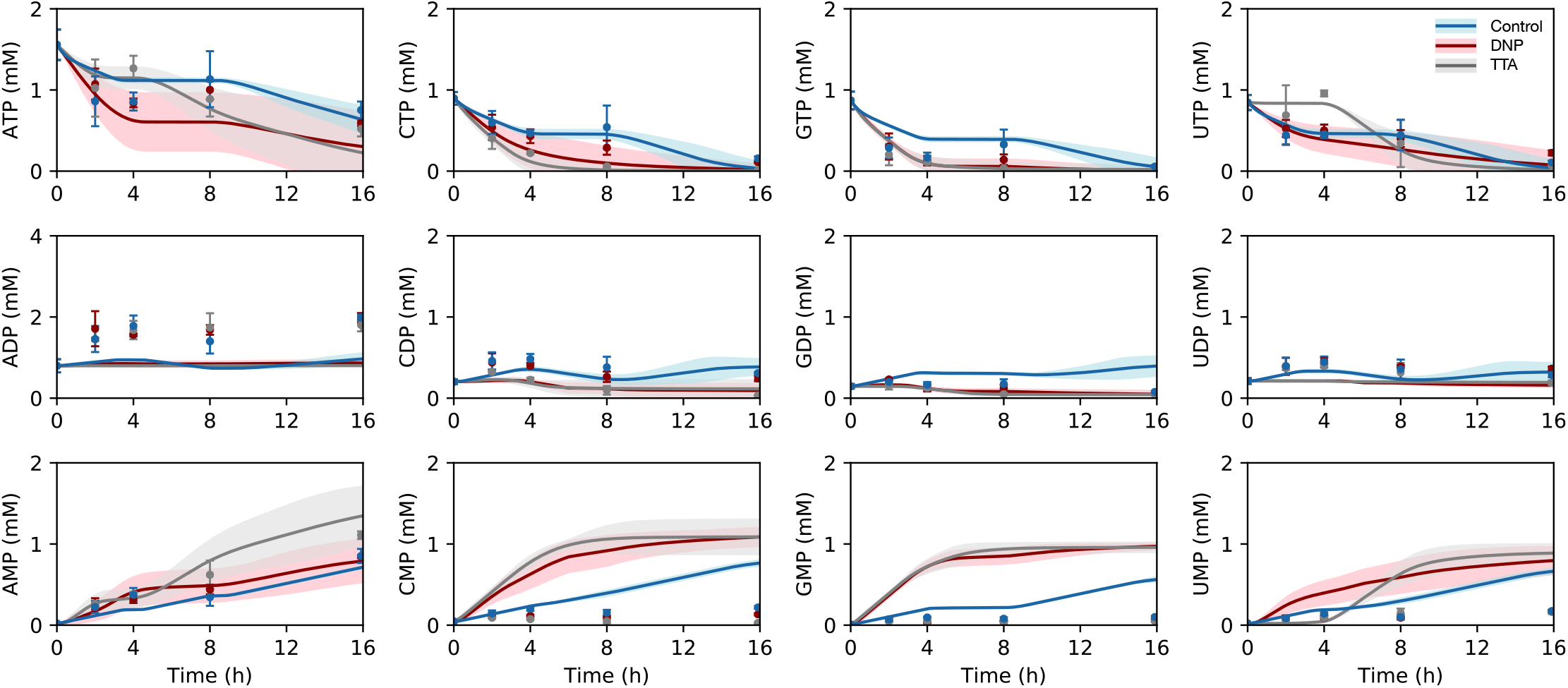
Time course of energy species levels in CFPS for control (blue), DNP (red), and TTA (grey). Both DNP and TTA exhausted GTP, which is required for translation, within 4 hours of the start of the reaction. Experimental fluxes constrained the mathematical model of CFPS. The solid line denotes the mean of the ensemble (N=100), the shaded region denotes the 95% confidence interval of the ensemble, the points denote experimental measurements, and error bars denote the standard deviation of experimental measurements.

**Fig. S6:**
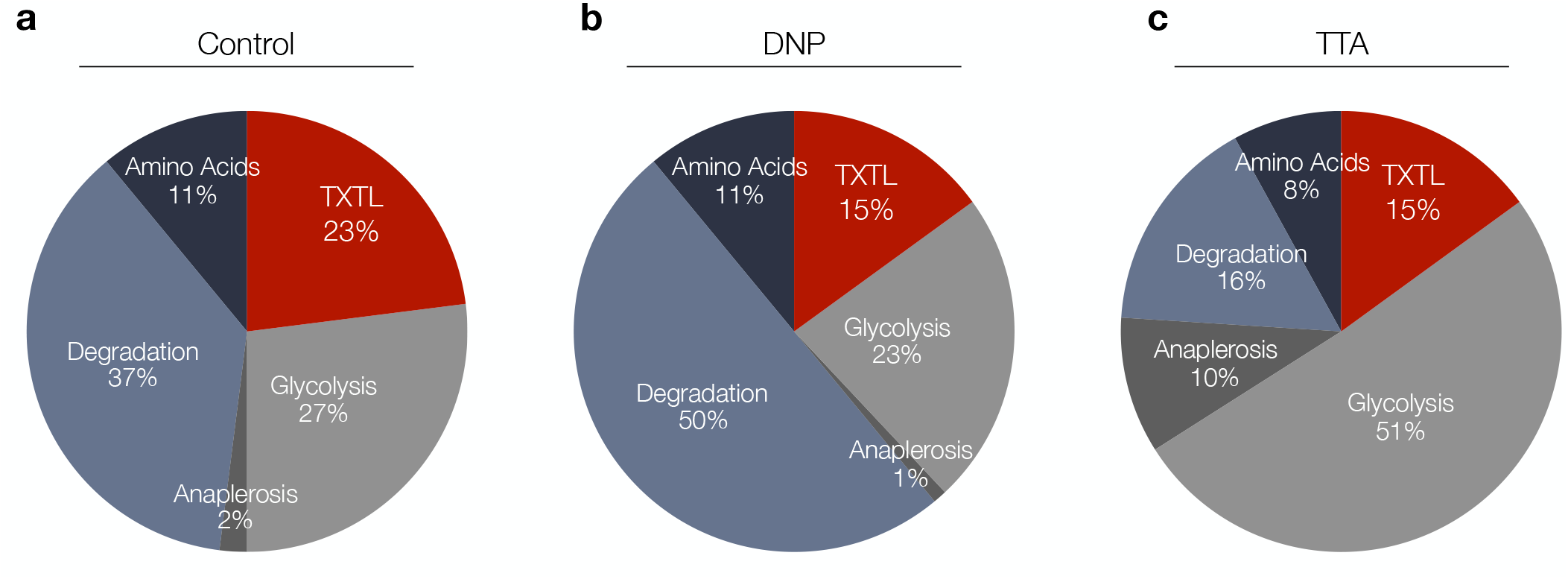
Mean energy efficiency across an ensemble (N=100) for control (a), DNP (b), and TTA (c) throughout the metabolic network. TXTL denotes the energy efficiency for transcription and translation processes.

**Fig. S7:**
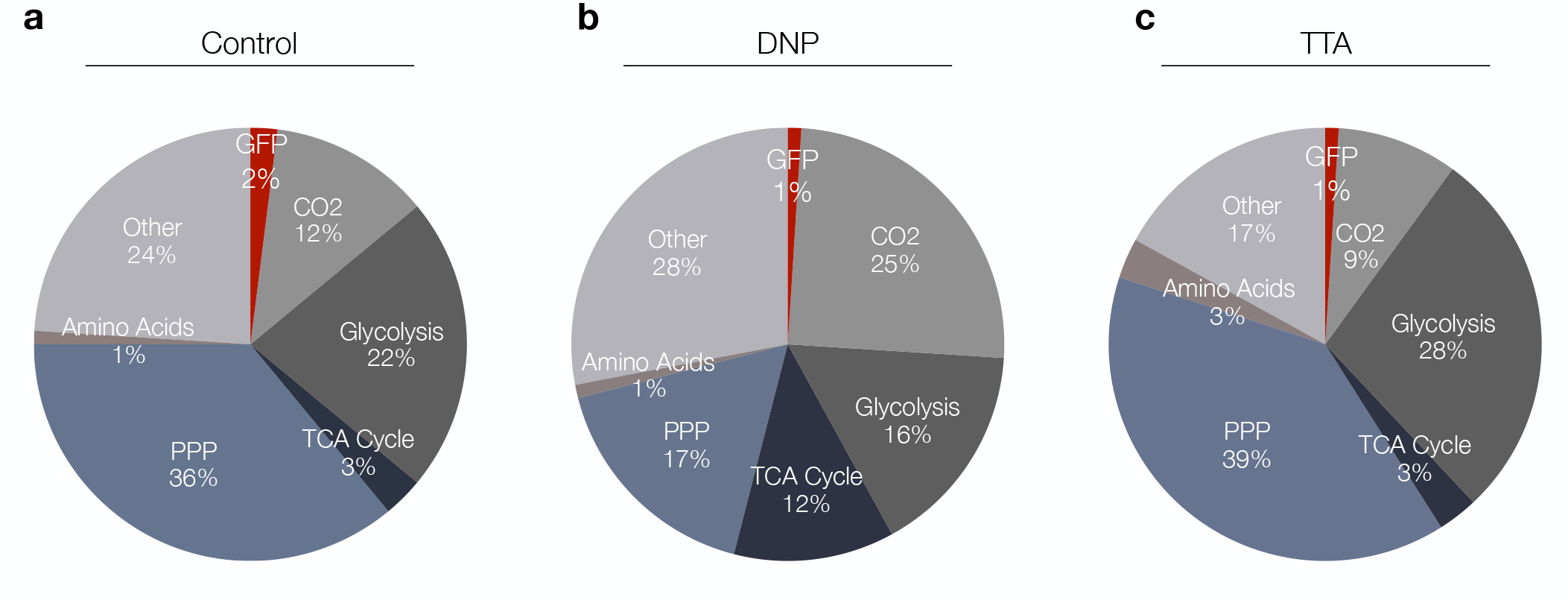
Mean carbon yield across an ensemble (N=100) for control (a), DNP (b), and TTA (c) for CFPS. PPP denotes the Pentose Phosphate Pathway. Other includes purine, pyrimidine and chorismate metabolism.

